# Engineering high-yield biopolymer secretion creates an extracellular protein matrix for living materials

**DOI:** 10.1101/2020.08.31.276303

**Authors:** Marimikel Charrier, Maria Teresa Orozco-Hidalgo, Nicholas Tjahjono, Dong Li, Sara Molinari, Kathleen R. Ryan, Paul D. Ashby, Behzad Rad, Caroline M. Ajo-Franklin

**Author notes:** These two authors contributed equally (Marimikel Charrier, Maria Teresa Orozco-Hidalgo).

## Abstract

The bacterial extracellular matrix forms autonomously, giving rise to complex material properties and multicellular behaviors. Synthetic matrix analogues can replicate these functions, but require exogenously added material or have limited programmability. Here we design a two-strain bacterial system that self-synthesizes and structures a synthetic extracellular matrix of proteins. We engineered *Caulobacter crescentus* to secrete an extracellular matrix protein composed of elastin-like polypeptide (ELP) hydrogel fused to Supercharged SpyCatcher (SC^(-)^). This biopolymer was secreted at levels of 60 mg/L, an unprecedented level of biopolymer secretion by a gram-negative bacterium. The ELP domain was swapped with either a crosslinkable variant of ELP or resilin-like polypeptide, demonstrating this system is flexible. The SC^(-)^-ELP matrix protein bound specifically and covalently to the cell surface of a *C. crescentus* strain that displays a high-density array of SpyTag peptides via its engineered Surface-layer. Our work develops protein design rules for Type I secretion in *C. crescentus*, and demonstrates the autonomous secretion and assembly of programmable extracellular protein matrices, offering a path forward towards the formation of cohesive engineered living materials.

**IMPORTANCE:** Engineered living materials (ELM) aim to mimic characteristics of natural occurring systems, bringing the benefits of self-healing, synthesis, autonomous assembly, and responsiveness to traditional materials. Previous research has shown the potential of replicating the bacterial extracellular matrix (ECM) to mimic biofilms. However, these efforts require energy intensive processing or have limited tunability. We propose a bacterially-synthesized system that manipulates the protein content of the ECM, allowing for programmable interactions and autonomous material formation. To achieve this, we engineered a two-strain system to secrete a synthetic extracellular protein matrix (sEPM). This work is a step towards understanding the necessary parameters to engineering living cells to autonomously construct ELMs.

## INTRODUCTION

Bacterial cells mold their environment through their extracellular matrix (ECM): a heterogeneous matrix of predominately polysaccharides with a mix of proteins, nucleic acids, and minerals (1). The autonomously-produced ECM is dynamic and bacteria vary its charge, hydrophobicity, porosity, or other properties to assist the cell with survival in various environments. Biofilm matrices function to facilitate mechanical stress tolerance (2), nutrient sorption, and both genetic and chemical communication (3–5). By interacting with the environment and controlling mass transfer, the matrix affects morphology, resilience, and interspecies interactions (3, 6) of the bacterial community, increasing its overall plasticity.

Engineered living materials (ELMs) attempt to mimic aspects of natural systems, including biofilms, and are poised to dramatically impact the fields of soft matter assembly and structural materials by adding abilities like self-healing, material synthesis, autonomous assembly, and responsiveness (7). Current synthetic biology tools (8), such as pioneering work with curli fibers (9–11) and bacterial cellulose (12, 13), modulate the endogenous ECM content but are limited in the sequence tunability of the biopolymer in the matrix and do not directly encapsulate individual bacteria, as an extracellular polysaccharide (EPS) layer does. Direct cell-encapsulation is largely approached using exogenous addition of polymers and attaching them through adhesive motifs or entrapment (14, 15). These approaches lack the autonomous formation of natural biofilms and thus require energy-intensive processing (15, 16) and added expense. Thus, there is an unmet need for a self-forming, yet programmable bacterial ECM.

Limited effort has been made to engineer the EPS layer, mostly because the confounding multistep syntheses of non-linear polysaccharides (17) make them difficult to program genetically. A more tractable approach to engineering the supramolecular structure of the ECM is to manipulate its protein content. We hypothesize that this simplification of the ECM to a synthetic extracellular protein matrix (sEPM) would result in more programmable interactions allowing for tunable 3D structures. Previous research shows that alterations in the composition of polypeptides with hydrogel-like behaviors, such as elastin or resilin, leads to different materials properties (18–20). The protein structure, degree of cross-linking, and number of weak interactions are all variables impacted by the peptide sequence (21, 22). In addition, protein-protein interactions can drive highly specific, selective, and even covalent binding, for example through the SpyCatcher-SpyTag system (23, 24).

The freshwater bacterium, *Caulobacter crescentus*, is emerging as a platform for synthetic biology and ELMs (25–29). This bacterium provides multiple advantages as a chassis: it is genetically tractable, is well-characterized due to its intriguing dimorphic life cycle (30), strongly adheres to surfaces via its holdfast matrix (31), and it has a modifiable proteinaceous Surface-layer (S-layer) (25, 32). In addition, it is oligotrophic and can flourish with minimal nutrients and in cold temperatures (33). We previously reported the construction of a set of *C. crescentus* variants (25), in which we engineered the S-layer protein, RsaA (34, 35), to display SpyTag, which is one part of the split-enzyme SpyTag/SpyCatcher system (36). These strains covalently ligate SpyCatcher-displaying inorganic nanocrystals, proteins, and biopolymers to the extracellular array at high density (25). With these advantages, *C. crescentus* is well positioned as a chassis for developing a sEPM.

To make the formation of a sEPM autonomous, high-level protein secretion is required. However, secretion of heterologous biopolymers has proven challenging for gram-negative bacteria and has not exceeded 30 mg/L (via the Type III secretion system) (37). While typically Type I secretion systems (T1SS) are considered to have low titers (38), the T1SS in *C. crescentus* has the potential to secrete high heterologous protein titers. The T1SS is endogenously tasked with transporting 10-12% of the total cell protein to form the RsaA surface-layer (39) and secretion of heterologous enzymes has been demonstrated (40). The T1SS is a one-step transport system that consists of an ABC transporter, membrane fusion protein, and outer membrane protein. The hallmark of T1SS substrates is the necessary C-terminal secretion signal. In addition, they typically include RTX domains with the nine residue consensus sequence GGxGxDxUx, wherein U is a hydrophobic residue, and these domains are usually involved in Ca^2+^ binding. As calcium is strictly regulated intracellularly at a level lower than required for these proteins to fold, it is presumed that TISS substrates remain largely unfolded until fully secreted (41). In *C. crescentus*, the RsaD-F_a,b_ T1SS transports the 1026 amino acid RsaA surface-layer protein (40). This system is unique in that it contains two homologous outer membrane proteins, RsaF_a_ and RsaF_b_ (42). The RsaA substrate protein contains an 82 amino acid secretion signal and six occurrences of the RTX domain. However, either the C-terminal 242 or 336 amino acids are required for maximal secretion of protein (39).

In this work, we put forth a new concept that employs an engineered two-strain system to create a bacterially-produced, sEPM that subsequently covalently coats the bacterial cell surface. We develop *C. crescentus* strains that use T1SS to export elastin-like polypeptides (ELP) or resilin-like polypeptides (RLP) fused to SpyCatcher^(-)^ at levels as high as 60 mg/L, twice the previously reported maximum for biopolymer secretion by a gram-negative bacterium (37). We then demonstrate the sEPM by binding purified SpyCatcher^(-)^-ELP fusion proteins covalently to our engineered SpyTag-displaying strain. Through our secretion efforts, we unravel design rules around folding and isoelectric point required to maximize biopolymer secretion via *C. crescentus*’s T1SS. Thus this work furthers understanding of Type I secretion and the value of *C. crescentus* as a secretion platform by demonstrating the self-synthesis and self-organization of a rationally designed and tunable synthetic extracellular protein matrix.

## RESULTS

### Design and Construction of *Caulobacter crescentus* Strains to Produce an Extracellular Matrix Protein

We sought to mimic key properties of naturally occurring biofilm matrices, while adding the structural and functional flexibility of synthetic ECMs to create a sEPM. This led to several molecular-level design constraints for our sEPM. First, to avoid the need to exogenously add materials, the matrix should be biologically synthesized and secreted extracellularly. Second, to mimic the structure and function of a natural ECM, with the potential for mechanical support, trapped hydration, and transport of nutrients and waste products, the matrix should be capable of forming a hydrogel. Third, to open the possibility for spatial patterning and to add robustness, the synthetic matrix should specifically and covalently bind cells of a different strain.

To meet these constraints, we designed a system consisting of two strains of *C. crescentus*: a ‘Secretor’ strain capable of synthesizing and secreting a proteinaceous extracellular matrix (**Fig 1a**) and a ‘Displayer’ strain capable of binding this protein on its surface at high density (**Fig 1b**). When co-cultured, we hypothesized that the Displayer strain would be coated in an autonomously formed sEPM, creating a living material (**Fig 1c**). Since we previously engineered versions of Displayer strains (25), our major task was to construct Secretor strains to secrete an extracellular matrix protein capable of specifically binding to Displayer strains.

**Figure 1:**
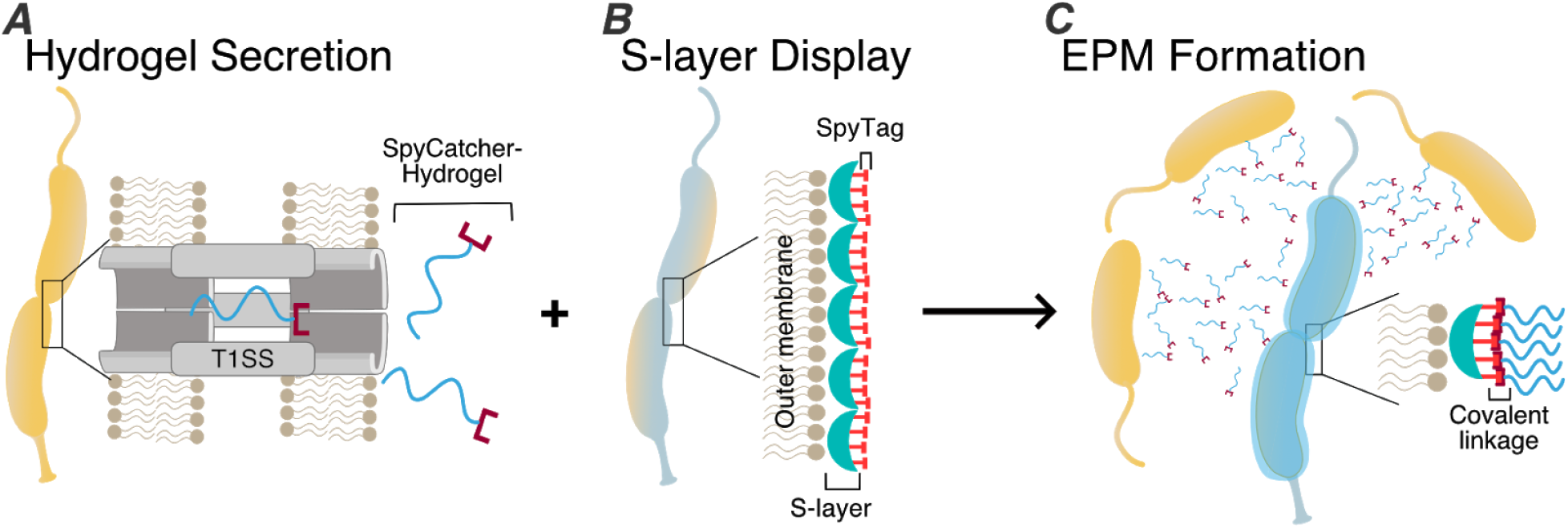
Design of two strain consortia that forms a synthetic extracellular protein matrix (EPM). Schematic of A) C. crescentus Secretor strain capable of exporting hydrogel proteins containing SpyCatcher covalent binding motif via a Type I Secretion System, B) C. crescentus Displayer strain capable of binding SpyCatcher-hydrogel proteins to engineered RsaA-SpyTag S-layer at high-density, and C) encapsulation of Displayer cells in the secreted hydrogel protein forming a synthetic EPM.

To create the Secretor strain, we designed a heterologous gene containing modules for high-level secretion, antibody-based detection, hydrogel formation, and covalent ligation (**Fig 2a**). This heterologous gene includes the native regulatory regions of *rsaA* (*CCNA_01059*), which drive high-level expression, and the C-terminal 336 amino acids of RsaA (notated 336c), which serve as a signal for extracellular secretion (43). While the 336c sequence can be detected via anti-RsaA antibodies (42), we also included an N-terminal FLAG tag so we could probe both termini. Since the Displayer strains display SpyTag (44), we utilized its partner, SpyCatcher, as the covalent binding motif. For the hydrogel module, we chose elastin-like polypeptide (ELP) which is a hydrophobic and disordered polymer that readily forms hydrogels (45). This thoroughly characterized recombinant protein has tunable material properties through alteration of the pentapeptide repeat number, inclusion of crosslinking residues, or inclusion of binding or cleavage domains (21, 46, 47). The ELP we utilize is nicknamed ELP_60_ due to its repeat number (SI Fig 1).

**Figure 2:**
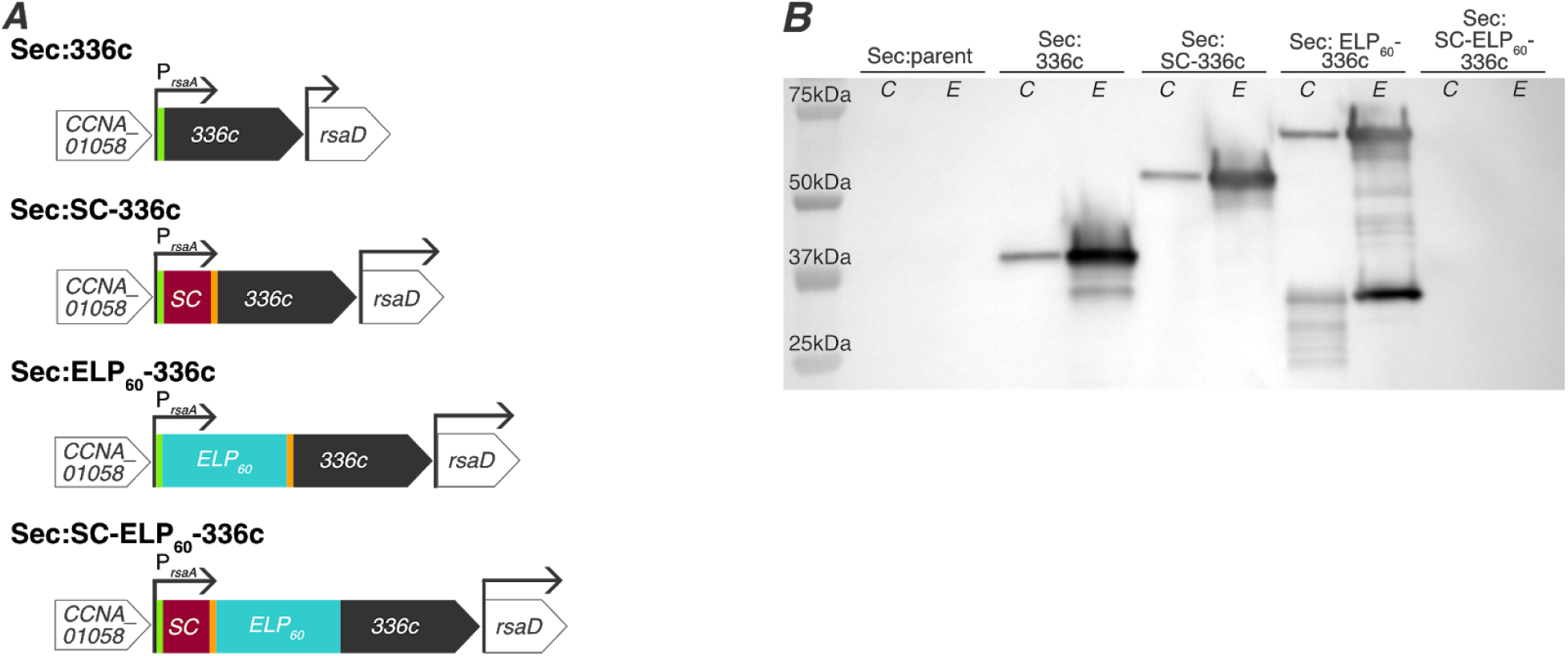
SpyCatcher-ELP60-336c unable to traverse T1SS in C. crescentus. A) Schematic of genome variants for secretion (top to bottom) of the 336c secretion signal, SpyCatcher(SC)-336c control, ELP_60_-hydrogel, and full SC-ELP_60_ protein. The green segment indicates a FLAG tag and orange segment indicates a Strep tag. All genes contain the native *rsaA* promoter and 5’ UTR. B) Immunoblot with anti-FLAG antibodies against whole cell lysate (indicated by C) and extracellular media (indicated by E) of Sec:parent (lanes 1c,1e) and Sec:336c (lanes 2c,2e), Sec:SC-336c (lanes 3c,3e), Sec:ELP_60_-336c (lanes 4c,4e), and Sec:SC-ELP_60_-336c (lanes 5c,5e). While full-length proteins are detected in the extracellular media in the strains expressing 336c, SC-336c, and ELP_60_-336c, the SC-ELP_60_-336c protein is not detected in either the cell pellet or extracellular media.

In addition to the heterologous gene that contained all modules, we also created genes with only the 336c signal, without the binding module, and/or without the hydrogel module as controls (**Fig 2A**). We introduced these DNA constructs into modified *C. crescentus* NA1000 strain (**Table 1**). NA1000 lacks the holdfast, making it less adherent to surfaces (48), and is a standard strain for studies of the *C. crescentus*. In our modified NA1000 strain, the native S-layer associated protein gene (*sapA*) is replaced with a xylose-inducible *mKate2 (49)* fluorescent protein gene to eliminate interference with S-layer assembly (50) and to facilitate fluorescent imaging. This parent strain is titled *Sec:parent*. All genome integrations were confirmed by colony PCR (SI Fig 2). The resultant Secretor strains are titled *Sec:SC-hydrogel variant* (genotype details available in SI Fig 1).

### Extracellular Matrix Proteins Containing a folded SpyCatcher Module are not Efficiently Secreted

To test the ability of our engineered strains to secrete synthetic extracellular matrix proteins, we analyzed the proteins present in the cell pellets and extracellular media using immunoblotting with an anti-FLAG antibody (**Fig 2B**). We did not detect any bands in the Sec:parent culture (**Fig 2**, lanes 1,2), confirming that our assay is specific for FLAG tag-containing proteins. In the Sec:336c culture, we observed a 37 kDa band corresponding to the 336c secretion signal. This band was notably stronger in the extracellular medium than the cell pellet (**Fig 2**, lanes 3,4), confirming earlier work that this sequence is sufficient for secretion (43). Similarly, somewhat weaker bands corresponding to SpyCatcher-336c (57 kDa) and ELP_60_-336c (73 kDa) were observed in extracellular media from the Sec:SC-336c and Sec:ELP_60_-336c cultures respectively (**Fig 2B**, lanes 5-8), indicating that these proteins were secreted. We noted degradation of ELP_60_-336c in both the cell pellet and supernatant fractions, with lower-molecular weight bands ranging from approximately 53 kDa to 20 kDa. Most critically, we were unable to detect the SpyCatcher-ELP_60_-336c fusion protein in either the supernatant or cell pellet (**Fig 2**, lanes 9,10), indicating that the complete synthetic extracellular matrix protein is either not secreted and subsequently degraded intracellularly or not expressed. Since the heterologous gene is present in the genome (SI Fig 2E), we suggest that the SpyCatcher-ELP_60_-336c fusion is unable to traverse the Type I machinery because of the folded nature of SpyCatcher (51, 52).

### Replacing SpyCatcher with Supercharged SpyCatcher^(-)^ Enables Secretion of Extracellular Matrix Proteins

To test the hypothesis that the folded nature of SpyCatcher decreases secretion of fusion proteins, we replaced SpyCatcher with a supercharged variant, SpyCatcher (SC^(-)^), in our synthetic extracellular matrix protein. SC^(-)^ has an additional 12 negative charges introduced into the sequence, which keeps it largely disordered until it binds SpyTag (44). We verified genomic incorporation of SC^(-)^ into the Sec:SC-ELP_60_-336c strain by colony PCR (SI Fig 1F) and notated it as Sec:SC^(-)^-ELP_60_-336c (**Fig 3A**).

**Figure 3:**
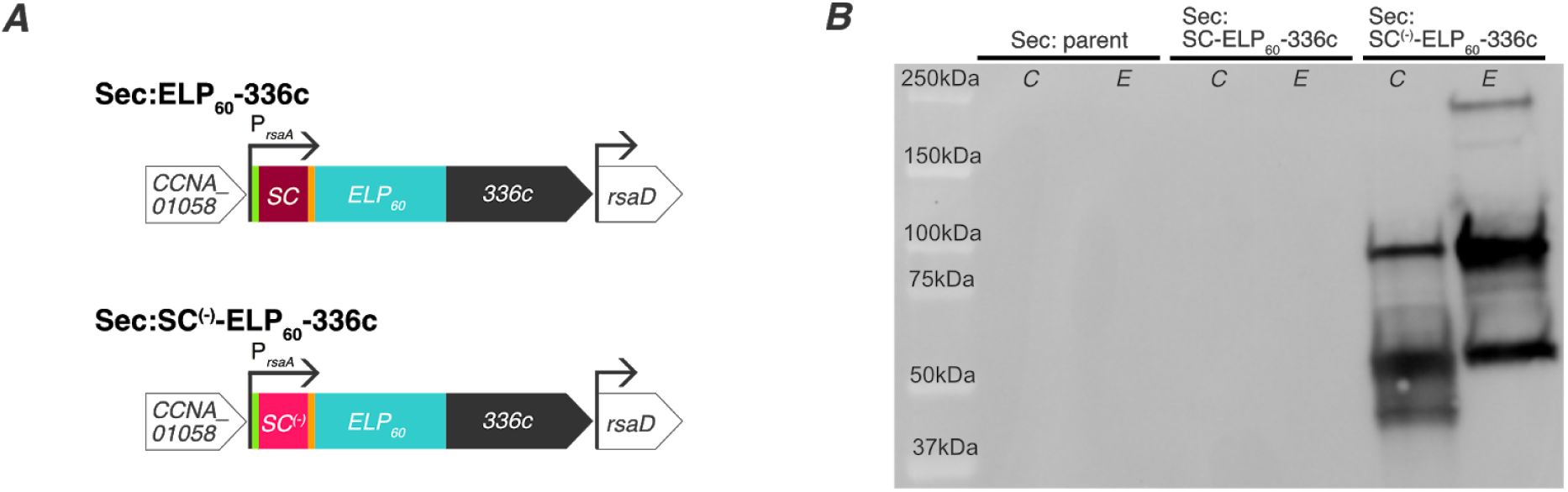
Supercharged SpyCatcher^(-)^ allows for secretion of synthetic extracellular matrix proteins. A) Schematic of genome variants for secretion of SpyCatcher-ELP_60_-336c protein and supercharged SpyCatcher(SC^(-)^)-ELP_60_-336c protein. The green segment indicates a FLAG tag and orange segment indicates a Strep tag. All genes contain the native rsaA promoter and 5’ UTR. B) Immunoblot with anti-FLAG antibodies against the whole cell lysate (indicated by C) and extracellular media (indicated by E) of Sec:parent (lanes 1c,1e), Sec:SC-ELP_60_-336c (lanes 2c,2e), and Sec:SC^(-)^ELP_60_-336c (lanes 3c,3e). While the matrix protein is not detected in the wild-type CB15 and Sec:SC-ELP_60_-336c cultures, the Sec:SC^(-)^-ELP60-336c strain achieves notable levels of secretion.

We analyzed the protein composition of the Sec:parent, Sec:SC-ELP_60_-336c, and Sec:SC^(-)^-ELP_60_-336c cultures by immunoblotting (**Fig 3B**). As expected, no FLAG tag-containing bands were present in either the cell pellet or supernatant fractions of the Sec:parent culture (lanes 1,2) or the Sec:SC-ELP_60_-336c strains (lanes 3,4). However, we detected a band at an apparent molecular weight of 93 kDa in the extracellular media of Sec:SC^(-)^-ELP_60_-336c cultures (lanes 5,6). While this apparent molecular weight is greater than the predicted molecular weight of SC^(-)^-ELP_60_-336c (86.9 kDa); previous work has demonstrated that SpyCatcher^(-)^ migrates as a ∼20 kDa protein, although its expected molecular weight is 14.8 kDa(44). Accounting for this difference, we expect SC^(-)^-ELP_60_-336c to run as a 92 kDa protein, in line with the observed band. This band is also detected with anti-RsaA serum, indicating that this band is indeed the full-length SC^(-)^-ELP_60_-336c (SI Fig 3). Hence, this confirms our hypothesis that replacing a folded fusion protein (Spycatcher) with a disordered one (SpyCatcher^(-)^) allows for expression and secretion due to the specificity for unfolded substrates by Type I secretion machinery.

There were significant bands at higher molecular weight (∼250 kDa) and lower molecular weight (∼52 and 41 kDa) present in the cell pellet and extracellular media of Sec:SC^(-)^-ELP_60_-336c cultures (**Fig 3B**, lanes 5,6). The higher molecular weight band became more pronounced as the protein concentration increased (SI Fig 4). As ELP_60_ is highly hydrophobic and the 336c secretion sequence tends to aggregate (43), so we attribute the 250 kDa band to aggregation. The presence of molecular weight bands lower than expected suggests that the protein is sensitive to degradation. Nevertheless, we detect most of the protein at the molecular weight expected for SC^(-)^-ELP_60_-336c, indicating this protein is effectively secreted.

**Figure 4:**
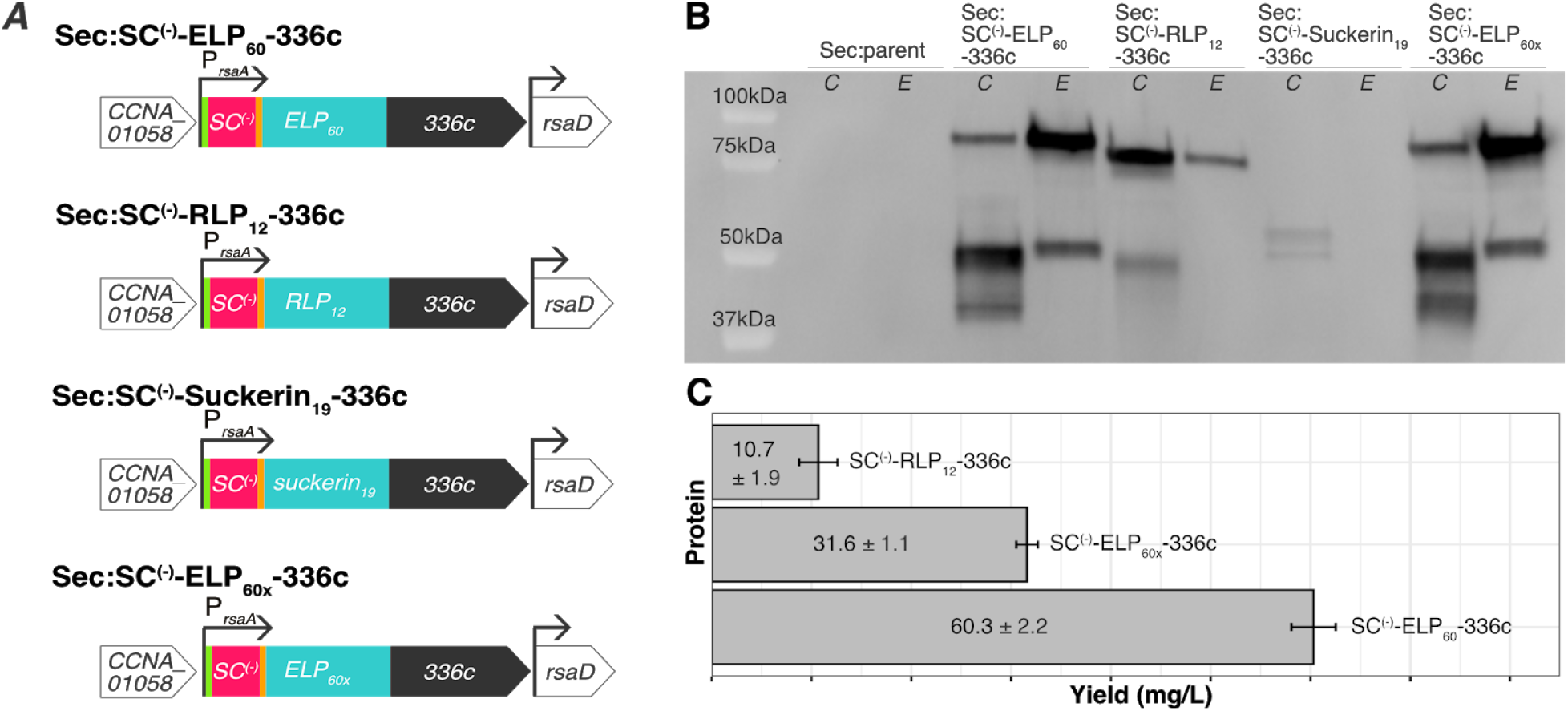
SC^(-)^ELP_60_ -336c, SC^(-)^ELP_60x_ -336c, and SC^(-)^RLP_12_ -336c are secreted, while SC^(-)^Suckerin_19_ -336c is not. A) Schematic of genome variants for secretion of (top to bottom) SpyCatcher^(-)^ELP_60_-336c protein, SpyCatcher^(-)^RLP_12_-336c protein, SpyCatcher^(-)^Suckerin_19_-336c protein, and SpyCatcher^(-)^ELP_60x_-336c protein. The green segment indicates a FLAG tag and orange segment indicates a Strep tag. All genes contain the native rsaA promoter and 5’ UTR. B) Immunoblot with anti-FLAG antibodies of *C. crescentus* whole cell lysate (indicated by C) and extracellular media (indicated by E) of Sec:parent (lanes 1c,1e), Sec:SC^(-)-^ELP_60_-336c (lanes 2c,2e), Sec:SC^(-)^RLP_12_-336c (lanes 3c,3e), Sec:SC^(-)^Suckerin_19_-336c (lanes 4c,4e), and Sec:SC^(-)^ELP_60x_-336c (lanes 5c,5e). While full-length proteins are detected in the supernatant in the strains expressing SC^(-)^ELP_60_-336c, SC^(-)^RLP_12_-336c, and SC^(-)^ELP_60x_-336c, the SC^(-)^Suckerin_19_-336c protein is only faintly apparent the cell pellet and not at all in the extracellular media. C) Average yield of protein purified from the extracellular media of Sec:SC^(-)^ELP_60_-336c cultures, Sec:SC^(-)^RLP_12_-336c cultures, and Sec:SC^(-)^ELP_60x_-336c cultures. Sec:SC^(-)^ELP_60_-336c is capable of exporting the highest quantity of a biopolymer by a bacteria to our knowledge. Values in the graph are represented as average protein yield in milligrams per liter of culture ±SD (n=3).

### Extracellular Matrix Proteins with Different Hydrogel Modules can be Secreted at High Levels

Next, we sought to test the modularity of the hydrogel component so that the material properties of sEPM could be tailored. We sought polypeptides that would cover a range of potential material properties while offering variety in hydrophobicity and structure (**Fig 4A**). We also required that our selected targets had previously been expressed heterologously and could tolerate sequence insertions.

Using these design criteria, we selected three additional hydrogel-forming polypeptides to incorporate into our sEPM. First, we created a variant of the ELP_60_ sequence, ELP_60x_, that includes a series of lysine and glutamine residues for enzymatic crosslinking with transglutaminase, potentially stiffening resultant ELMs (53). Our second target, resilin-like polypeptide (RLP_12_) (19), is a recombinant version of an elastomeric protein found in insects that has remarkable extensibility and resilience. While less explored than ELP, several variants of RLP have been produced with modular sequences for tunable material properties, including sites for enzymatic crosslinking and inclusion of biologically active domains (20). Similar to ELP_60_, RLP_12_ is disordered. However RLP_12_ is more hydrophilic than ELP_60_. Finally, we selected Suckerin_19_, a protein found in the sucker ring teeth of squid and cuttlefish. This unique material is highly stiff, with a potential elastic modulus in the gigapascal range (54). It differs from the two other targets not only in stiffness, but also in that it is a structured protein of beta-sheets interspersed with amorphous regions. These three new heterologous genes were inserted into the genome in place of the first ELP_60_ gene to create the Sec:SC^(-)^-ELP_60x_-336c, Sec:SC^(-)^-RLP_12_-336c, Sec:SC^(-)^-Suckerin_19_-336c strains, and all were confirmed by colony PCR (SI Fig 1).

Again, we used immunoblotting to identify proteins from these engineered Secretor strains. The Sec:SC^(-)^-ELP_60_-336c and Sec:SC^(-)^-ELP_60x_-336c cultures displayed bands of 90 kDa in the extracellular media (**Fig 4B**, lanes 3,4 and 9,10 respectively), demonstrating that ELP_60_ sequence can be tuned while maintaining secretion. Additionally, a weaker band of 81 kDa corresponding to SC^(-)^-RLP_12_ was detected in the supernatant (**Fig 4B**, lanes 5,6), indicating it also can be secreted. However, the expected band for the SC^(-)^-Suckerin_19_ protein (97.9 kDa) was not detected, but degradation products at lower molecular weight bands (53 kDa and 46kDa, **Fig 4B**, lanes 7,8) were apparent in the cell pellet at very low levels. Taken together, these results confirm that synthetic extracellular matrix proteins containing different hydrogel polypeptides can be secreted, but that the hydrogel protein sequence has a significant effect on secretion yields.

To determine the total amount of extracellular matrix protein secreted by the engineered strains, we purified the protein from culture media using ion exchange chromatography (SI Fig 5) and measured concentration using a bicinchoninic acid (BCA) assay. The Sec:SC^(-)^-ELP_60_ -336c strain secreted the highest amounts of protein, 60.3 ± 2.22 mg of SC^(-)^-ELP_60_ -336c per liter of culture (Fig 4C). Interestingly, the Sec:SC^(-)^-ELP_60x_ -336c strain secreted roughly half as much as the ELP_60_ variant (31.6 ±1.07 mg/L culture, p<0.001), and Sec:SC^(-)^-RLP_12_ -336c secreted approximately one-sixth as much protein (10.7 ±1.91 mg/L culture, p<0.0001). Significant differences in yield persist even when these values are normalized to the number of cells in the culture (SI Fig 6). Since SC^(-)^-ELP_60_ variants and SC^(-)^-RLP_12_ are predicted to have a disordered structure while SC^(-)^-Suckerin_19_ is predicted to be structured, these results support the hypothesis that secretion through T1SS is strongly affected by protein structure. Moreover, the high yields of secreted protein support *C. crescentus* as a multi-functional ELM chassis.

**Figure 5:**
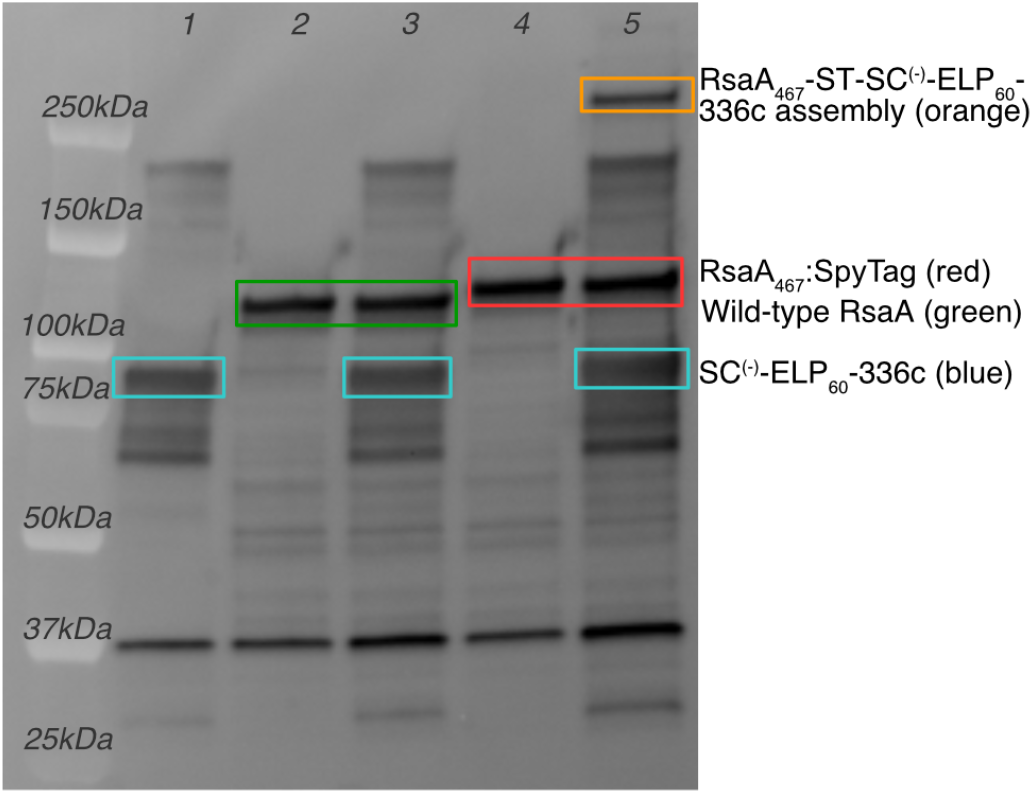
Purified SC(-)-ELP60-336c can be covalently bound to RsaA467-ST on Displayer cells. A high-molecular weight band corresponding to the covalent assembly of purified SpyCatcher^(-)^-ELP_60_-336c protein and RsaA_467_-SpyTag on engineered Disp:RsaA_467_-ST cells with is detected with anti-RsaA antibodies (lane 5, orange box). This band is not detected in control lanes containing only purified SC^(-)^-ELP_60_-336c protein (lane 1), only Disp:RsaA_wt_ cells (lane 2), purified SC^(-)^-ELP_60_-336c protein with Disp:RsaA_wt_ cells (lane 3), or only engineered Disp:RsaA_467_-ST cells (lane 4).

### Design and Construction of a Strain for Covalent Binding of Extracellular Matrix Proteins

We previously reported a *C. crescentus* NA1000 strain capable of displaying SpyTag on the cell surface via insertion into the RsaA S-layer lattice (25). The variant with the highest level of SpyCatcher-mRFP1 surface binding, over 11,000 copies per cell, contains SpyTag at amino acid location 467 of RsaA. For this study, we utilized the same SpyTag insertion but in the CB15 background, as opposed to the NA1000 background. CB15 is desirable for material formation as it retains the holdfast protein matrix at the end of the stalk structure, which allows *C. crescentus* stalked cells to adhere strongly to surfaces. Our rationale for using this background, as opposed to NA1000, is that this is an additional point of control contributing to the material properties as it could allow for strong attachment of inorganic particles to our system. As before, we engineered a fluorescent protein into the CB15 background strain, this time GFPmut3 (55), in place of SapA which could undesirably edit our modified S-layer proteins. The final experimental construct is titled Disp:RsaA_467_-ST and a control strain that does not contain the SpyTag modification is titled Disp:RsaA_wt_(genotype details available in SI). The SpyTag insertion was confirmed by colony PCR (SI Fig 1J).

### Extracellular Matrix Protein Binds Specifically and Covalently to Create a sEPM

Next, we sought to test whether the synthetic extracellular matrix proteins were capable of ligation to SpyTag on the Displayer strain, creating a sEPM. We incubated the Disp:RsaA_467_-SpyTag with purified SC^(-)^-ELP_60_-336c protein and analyzed the reaction by immunoblotting with anti-RsaA polyclonal antibodies (**Fig 5**). In control samples containing only SC^(-)^-ELP_60_-336c protein (**Fig 5**, lane 1, 83 kDa band), Disp:RsaA_wt_ alone (**Fig 5**, lane 2, 112 kDa band), SC^(-)^-ELP_60_-336c protein with Disp:RsaA_wt_(**Fig 5**, lane 3, 83 kDa and 113 kDa bands respectively), or Disp:RsaA_467_-ST cells alone (**Fig 5**, lane 4, 121 kDa band), bands at 83kDa or 112kDa were detected since all samples contain some of RsaA, either the full RsaA protein (Displayer cells) or the C-terminal 336c uncleaved secretion signal (purified protein). We note the apparent molecular weight of the purified SC^(-)^-ELP_60_ -336c protein is approximately 10 kDa lower than seen in secreted protein prior to purification. We attribute this change in gel migration pattern to charge screening of these proteins by the PBS solution, which may decrease aggregation and/or alter protein conformation (56, 57). A higher molecular weight band associated with covalent bonding between SC^(-)^-ELP_60_ -336c and RsaA_467_ -ST in the control samples was not observed, which is consistent with the requirement for both SpyTag and SpyCatcher^(-)^ to be present for ligation to occur. In the sample containing both SC^(-)^-ELP_60_ -336c protein and Disp:RsaA_467_-ST cells, a higher molecular weight product is apparent (**Fig 4**, lane 5). These observations indicate that a specific covalent attachment is formed between the fusion protein and the Disp:RsaA_467_-ST cell’s engineered RsaA S-layer through the SpyCatcher^(-)^-SpyTag system, resulting in hydrogel coating of the Displayer strain and formation of a sEPM.

## DISCUSSION

As demonstrated above, we constructed a modular extracellular protein matrix through secretion of hydrogel materials that covalently coat cells. A switch to the supercharged SpyCatcher^(-)^ variant enables the extracellular matrix proteins to be secreted via a T1SS at unprecedented levels. Extracellular matrix proteins with hydrogel domains of elastin-like polypeptide (ELP_60_ and ELP_60x_) or a resilin-like polypeptide (RLP_12_) can be secreted, demonstrating the modularity of our approach. The extracellular matrix protein binds specifically to our engineered RsaA_467_-SpyTag S-layer, enveloping the outermost cell surface and creating a sEPM. In the following, we discuss how our findings impact our understanding of Type I secretion in *C. crescentus*, and new routes towards self-coating bacteria and autonomous assembly of engineered living materials.

### Engineered *C. crescentus* is a platform for high-level secretion of biopolymers

This work achieved unprecedented levels of biopolymer secretion in Gram negative bacterial hosts, doubling previously reported levels (37). In our research, we discovered that through the switch of Spycatcher to SpyCatcher^(-)^, we achieve secretion of heterologous polymer-protein fusions (**Fig 3B**), accomplishing the highest reported yields (60.3 ± 2.22 mg/L) of a secreted biopolymer (SC^(-)^-ELP_60_ -336c) by a Gram-negative bacterium (38) (**Fig 4B**). We hypothesize that SpyCatcher^(-)^ fusions are required in this system because SpyCatcher^(-)^ remains largely disordered until it partners with SpyTag and the T1SS machinery has a strong preference for unfolded proteins. This hypothesis is further supported by the fact that we are unable to secrete fusions involving the Suckerin_19_ protein, as it contains structured beta sheets (51, 52, 58).

Moreover, our observations that the different polymer-protein fusions vary in secretion yields (**Fig 4C**) uncovers design strategies for maximizing heterologous protein secretion through *C. crescentus*’s T1SS. For instance, SC^(-)^-RLP_12_ -336c is secreted at significantly lower levels than SC^(-)^-ELP_60_ -336c. Previous studies have shown that ABC transport systems, such as the RsaD-F_a,b_ T1SS used herein, have higher secretion yields with proteins with pI’s lower than 5.5. This pI selectivity is ascribed to the conformational changes of the transport machinery when it interacts with the target protein and the electric potential of cell membranes (59). While all of the successfully secreted proteins have an overall pI lower than 5.5, the lowest pI being 3.83 for 336c and the highest pI being 5.09 for SC^(-)^-ELP_60x_-336c, the pI’s of the hydrogel domains within the full-length proteins vary greatly (SI Fig 7). Thus, we attribute the robust secretion yield of SC^(-)^-ELP_60_ -336c to the ELP_60_ domain’s pI of 5.5, and the low secretion yield of SC^(-)^-RLP_12_ -336c to the high isoelectric point (pI) of 9.91 for the RLP_12_ domain. The ELP_60x_ domain also has a high pI of 10.70, and accordingly secretion levels of SC^(-)^-ELP_60_ -336c are lower than that of SC^(-)^-ELP_60x_ -336c (**Fig 4C**). We also postulate that SpyCatcher was able to be secreted (**Fig 2B**) despite that it is a folded protein because of its low pI of 4.14. This result confirms previous work showing that secretion of ELPs is affected by amino acid sequence, credited to the shift in surface chemistry interactions (60). Overall, our work sheds light on key design criteria for high yield secretion targets using *C. crescentus* T1SS. To be secreted at high yield, the protein should have i) minimal, ideally no, regions with secondary or tertiary structure, ii) an overall pI lower than 5.5, and iii) individual domains with pIs lower than 5.5.

Our work indicates secreted biopolymers can easily be purified from *C. crescentus* cultures through anion exchange chromatography of the extracellular media, without the need for cell lysis. Since *C. crescentus* secretes few extracellular proteins, there are fewer contaminants to remove from the target protein. These advantages are beneficial for applications where extremely pure hydrogel material is desired without expensive processing, establishing *C. crescentus* as a powerful chassis for secretion of different biopolymers.

### sEPM-forming consortia have applications in biomanufacturing and engineered living materials

We found that the SC^(-)^-ELP_60_ -336c biopolymer binds covalently to the cell surface of our *C. crescentus* Displayer strain via an engineered S-layer array. While there are many reports of cell encapsulation in chemically produced hydrogels (14, 16, 61–63), this is the first report of entirely bacterial synthesized, covalent layering of hydrogel material on a bacterial cell surface. The use of spontaneously aggregating polymers like ELP and RLP advances our previously reported 2D assembly of a biomaterial onto a cell surface (25) into a 3D material. Looking forward, we envision the ability to engineer self-synthesis and self-organization of a rationally-designed synthetic extracellular protein matrix will lead to autonomously assembled living materials. Such a material could be formed without intervention, induction chemicals, or applied force via an oligotroph that does not require complex nutrients due to its high synthetic capacity (33, 64), making it a uniquely low-cost, low-effort living material. These ELMs would also be modular based on the biopolymer design and cell count. This platform could be applied to streamlined biomanufacturing of complex materials but also chemicals or fuels, as enveloping cells in a hydrogel can assist with cell protection in large-scale production.

In addition to self-encapsulation for biomanufacturing, we suggest this approach has additional benefits for ELMs. First, by localizing the matrix protein to the cell surface, a hydrogel-like biomaterial may assemble under lower solution-phase protein concentrations than is usually reported for creation of a strong hydrogel. Second, this Displayer strain has been engineered in a background strain of *C. crescentus* that still retains the holdfast matrix at the base of its stalk, as opposed to the Secretor strain where this holdfast is no longer present. This allows for Displayer cells to adhere strongly to surfaces and could be used to integrate inorganic materials into a hybrid material, such as introduction of orthogonal mechanical or optoelectronic properties (14, 65). The holdfast also opens the door to cell patterning through chemical modification of the surface (66) or layer-by-layer deposition of the ELM through bioprinting (67–69). This hierarchical assembly and cell patterning can lead to mechanical properties found in natural systems such as tolerance of compressive force (70) or arresting crack propagation (71, 72). Third, since the sEPM is self-synthesized, damage to the hydrogel layer can be continually repaired, the material can expand over time, and a small sample of the ELM can nucleate the growth of more material. Fourth, *C. crescentus* is non-pathogenic, has lower endotoxin activity then *E. coli (73)*, and has been previously developed as a microbicide (27), making it a safe option for deployment. Upon further engineering control over the cell-patterning within the consortia, we also envision usage for this advanced material in self-healing infrastructure, soft robotics, bioremediation, and biomedicine.

In summary, we describe the creation of strains that secrete a synthetic extracellular protein matrix and demonstrate the cell-surface attachment of the sEPM. In doing so, we elucidated several design criteria for maximal biopolymer secretion via T1SS, including encoding an isoelectric point ≤ 5.5 for each domain and the entire protein, and limiting folded domains. Similar to a naturally occurring biofilm matrix, our engineered matrix is both composed of hydrogel-forming biomolecules and encodes specific binding to engineered strains. However, our engineered matrix binds through covalent bonds rather than weak interactions (3) and the matrix composition can potentially be altered to provide emergent properties. This work further develops *C. crescentus* as a chassis for high-level secretion, demonstrates a sEPM, and takes an important step forward towards creating autonomous, ordered ELMs for use in biomanufacturing and advanced materials.

## MATERIALS AND METHODS

### Strains

All strains used in this study are listed in Table S1. The *C. crescentus* strains were grown in PYE media (0.2% peptone, 0.1% yeast extract, 1 mM MgSO_4_, 0.5 mM CaCl_2_) at 20°C or 30°C and with aeration at 250 rpm. The *E. coli* strains were grown in LB media (1% tryptone, 0.5% yeast extract, 1%NaCl) at 37°C with aeration at 250 rpm. Depending on the strain used, antibiotics were used in the following concentrations: For *E. coli*, 50 μg/mL ampicillin and 30 μg/mL kanamycin. For *C. crescentus* 25 μg/mL kanamycin (plate). For conjugation methods, 300 μM diaminopimelic acid (DAP) was supplemented. For recombination methods, 3% w/v sucrose was supplemented. All chemicals were purchased from Sigma-Aldrich or VWR.

### Plasmid construction

The list of all plasmids and primers used in the present study is available in Tables S2-S3. Details on construction of pNPTS138 integration plasmids can be found in Supporting Information. Plasmids were introduced to *E. coli* using standard transformation techniques with NEB 5-Alpha chemically competent cells (New England BioLabs), and to *C. crescentus* using conjugation via *E. coli* strain WM3064.

### Genome Engineering of *C. crescentus*

In order to make the background strain (*C. crescentus* CB15N Δ*sapA*::Pxyl-*mkate2*), the *sapA* gene (*CCNA_00783*) was replaced with gene for the mKate2 fluorescent protein under a xylose induction promoter. In order to achieve this, the 2-step recombination technique with sucrose counterselection method was employed. The fusion gene sequences for all synthetic extracellular matrix proteins and control proteins were integrated into the genome in place of the native *rsaA* gene using a 2-step recombination technique with sucrose counterselection, leaving the native regulatory sequence intact.

The 2-step recombination technique with sucrose counterselection is as follows: the pNPTS138 plasmids were electroporated into *E. coli* WM3064 cells and subsequently conjugated overnight into *C. crescentus* CB15N Δ*sapA*::Pxyl-*mkate2* on a PYE agar plate containing 300μM DAP. The culture was then plated on PYE with kanamycin to select for integration of the plasmid and removal of WM3064 cells. Successful integrants were incubated in liquid PYE media overnight and plated on PYE supplemented with 3% (w/v) sucrose to select for excision of the plasmid and *sacB* gene, leaving the target sequence in the genome. Integration of the sequences and removal of *sacB* gene was confirmed by colony PCR (SI Fig 2) with OneTaq Hot Start Quick-Load 2X Master Mix with GC buffer (New England BioLabs) using a Touchdown thermocycling protocol with an annealing temperature ranging from 72-62°C, decreasing 1°C per cycle. Primer sets for colony PCR verification can be found in SI Table 3.

Protein expression and secretion was evaluated in cultures of 25 ml of PYE media with 0.02% Antifoam 204 (Sigma-Aldrich) after a 16 hr incubation at 20°C with aeration at 250 rpm from a starting OD_600_ of 0.02. Protein levels were evaluated through analysis of the whole cell lysate and extracellular media after separation by centrifugation at 10,000 RCF for 10 minutes. Fractions were combined with an equivalent volume of 4x Laemmli Buffer (BioRad) and incubated for 20 min at 95°C then run on a BioRad Criterion Stain-free 4-20% SDS-PAGE gel. The SDS-PAGE gel was transferred to a nitrocellulose membrane using the Bio-Rad Trans-Blot Turbo system, and blocked with Thermo-Fisher SuperBlock buffer for 1 hour with agitation. The membrane was then washed several times with Tris-Buffered Saline with 0.1% Tween 20 (TBST) before a 30 in incubation with monoclonal Mouse Anti-FLAG M2-Peroxidase (HRP) antibodies (Sigma-Aldrich) diluted 1:5000 in TBST. Thermo-Fisher SuperSignal West Pico Chemiluminescent Substrate was used to activate HRP fluorescence, and the membrane was imaged under Chemiluminescent mode in a ProteinSimple FluorChem E System. For SDS-PAGE gels, BioRad Precision Plus Protein Standards Unstained (Bio-Rad) was used. For immunoblot membranes, BioRad Precision Plus Protein Kaleidoscope Prestained Protein Standards (Bio-Rad) was used. Molecular weight band quantification was made using GelAnalyzer 19.1 software.

### Protein Quantification

To quantify protein secretion, engineered *C*.*crescentus* strains were cultured for 24 hours in PYE with 0.02% Antifoam 204 at 30°C with aeration at 250 rpm from a starting OD_600_ of 0.02. 25 ml cultures were used for Sec:SC^(-)^-RLP_12_ -336c and 250 ml cultures for Sec:SC^(-)^-ELP_60_ -336c and Sec:SC^(-)^-ELP_60x_ -336c.

After incubation, the cultures were centrifuged for 20 minutes (8000 RCF for 250 ml cultures, 5250 RCF for 25 ml cultures) to extract the extracellular solution. The extracellular media was then diluted two-fold with 20 mM HEPES buffer at pH 7.0 and applied to 1 column volume (CV) of DEAE Sepharose Fast Flow Resin (from GE Healthcare) equilibrated with 10 CV of 20 mM HEPES buffer at pH 7.0 (1 CV equaled 2 ml of settled resin for 25 ml cultures and 1 CV equaled 4 ml of settled resin for 250 ml cultures). The supernatant was allowed to flow-through the resin by gravity. The resin was subsequently washed with 10 CV of 20 mM HEPES buffer containing 50 mM NaCl at pH 7.0 and the protein was eluted with 3 CV of 20 mM HEPES buffer containing 500 mM NaCl at pH 7.0. This eluted fraction was placed in 12-14 kDa Molecular Weight Cut-off regenerated cellulose dialysis tubing (Spectra/Por) and dialyzed in phosphate-buffered saline (PBS) overnight at 4°C with stirring.

Protein quantification was performed using the protocol and supplies provided by the BCA Protein Assay Kit (Thermo Scientific Pierce). Samples of BSA protein in PBS ranging in concentration from 25 ug/ml to 2000 ug/ml were used to create the standard concentration curve. Triplicate measurements were performed for each sample to ensure consistent concentration readings. The determined concentration value was then multiplied by the total sample volume to obtain the protein yield value for the culture, and this value used to extrapolate the equivalent protein yield from a liter culture. Statistical significance was determined by an unpaired, two-tailed student’s t-test on Rstudios.

### Supercharged SpyCatcher-SpyTag Binding Assay

Displayer strains containing either *rsaA*_*wt*_ or *rsaA*_*467*_:*spytag* genes were grown at 30°C until they reached mid-log phase (OD_600_ 0.2-0.4). 10^8^ cells were harvested from each culture (as determined through optical density readings, where 1ml of OD_600_ =0.5 is equivalent to 10^9^ cells) and resuspended in 100 µl of PBS and 0.5 mM CaCl_2_. Purified SC^(-)^-ELP_60_ -336c protein was then added to either Disp:RsaA_467_-ST cells or Disp:RsaA_wt_ cells at a ratio of 1 RsaA monomer (74) to 50 SC^(-)^-ELP_60_ -336c monomers assuming 45,000 RsaA monomers per cell. In addition, three other negative controls were tested: Disp:RsaA_wt_ cells without protein, Disp:RsaA_467_-ST cells without protein, and SC^(-)^-ELP_60_ -336c protein without cells. All five of these mixtures were incubated for 3 days at 4°C with agitation, as this temperature was previously shown to increase SpyCatcher^(-)^ reactivity towards SpyTag (44). Afterwards, 15 ul from each sample was combined with an equivalent volume of 2x Laemmli Buffer (Bio-Rad) and boiled at 98°C for 20 minutes. The samples were analyzed via immunoblot as above, except using polyclonal Rabbit-Anti-C terminal RsaA antibodies (courtesy of the Smit lab, UBC (42)) diluted 1:5000 in TBST followed by Goat Anti-Rabbit IgG antibodies (HRP-conjugate, Sigma-Aldrich) diluted 1:5000 in TBST.

## Data availability

This study includes no data deposited in external repositories.

## ACKNOWLEDGEMENTS

We are indebted to Prof. John Smit and Dr. John Nomellini for helpful conversations and starting materials. This work was supported by the Defense Advanced Research Projects Agency (Engineered Living Materials Program, C.M.A-F.). Work at the Molecular Foundry was supported by the Office of Science, Office of Basic Energy Sciences, of the U.S. Department of Energy under Contract No. DE-AC02-05CH11231.

## AUTHOR CONTRIBUTIONS

M.C., B.R., and C.M.A.-F. contributed to Conceptualization. M.C., M.T.O.H., and N.T. contributed to Investigation, Methodology and Visualization. D. L. and S.M. contributed to Investigation and Methodology. M.T.O.H., D.L., and S.M. contributed Validation. K.R.R. and B.R. contributed Resources. M.C., K.R.R., P.D.A., B.R. contributed to Project Administration and Supervision. P.D.A. and C.M.A.-F. contributed to Funding Acquisition. M.C, M.T.O.H., N.T. and C.M.A-F. contributed to Writing – original draft. All authors contributed to Writing – review and editing.

## REFERENCES

1. Dragoš A, Kovács ÁT. 2017. The Peculiar Functions of the Bacterial Extracellular Matrix. Trends Microbiol 25:257–266.

2. Shaw T, Winston M, Rupp CJ, Klapper I, Stoodley P. 2004. Commonality of elastic relaxation times in biofilms. Phys Rev Lett 93:098102.

3. Flemming H-C, Wingender J. 2010. The biofilm matrix. Nat Rev Microbiol 8:623–633.

4. Sutherland I. 2001. Biofilm exopolysaccharides: a strong and sticky framework. Microbiology 147:3–9.

5. Steinberg N, Kolodkin-Gal I. 2015. The Matrix Reloaded: Probing the Extracellular Matrix Synchronizes Bacterial Communities. J Bacteriol 197:2092–2103.

6. Persat A, Nadell CD, Kim MK, Ingremeau F, Siryaporn A, Drescher K, Wingreen NS, Bassler BL, Gitai Z, Stone HA. 2015. The mechanical world of bacteria. Cell 161:988–997.

7. Nguyen PQ, Courchesne N-MD, Duraj-Thatte A, Praveschotinunt P, Joshi NS. 2018. Engineered Living Materials: Prospects and Challenges for Using Biological Systems to Direct the Assembly of Smart Materials. Adv Mater 30:e1704847.

8. Gilbert C, Ellis T. 2018. Biological Engineered Living Materials: Growing Functional Materials with Genetically Programmable Properties. ACS synthetic biology. ACS Publications.

9. Duraj-Thatte AM, Courchesne ND, Praveschotinunt P, Rutledge J, Lee Y, Karp JM, Joshi NS. 2019. Genetically Programmable Self-Regenerating Bacterial Hydrogels. Adv Mater 31:1901826.

10. Chen AY, Deng Z, Billings AN, Seker UOS, Lu MY, Citorik RJ, Zakeri B, Lu TK. 2014. Synthesis and patterning of tunable multiscale materials with engineered cells. Nat Mater 13:515.

11. Huang J, Liu S, Zhang C, Wang X, Pu J, Ba F, Xue S, Ye H, Zhao T, Li K, Wang Y, Zhang J, Wang L, Fan C, Lu TK, Zhong C. 2019. Programmable and printable Bacillus subtilis biofilms as engineered living materials. Nat Chem Biol 15:34–41.

12. Zogaj X, Nimtz M, Rohde M, Bokranz W, Römling U. 2001. The multicellular morphotypes of Salmonella typhimurium and Escherichia coli produce cellulose as the second component of the extracellular matrix. Mol Microbiol 39:1452–1463.

13. Florea M, Hagemann H, Santosa G, Abbott J, Micklem CN, Spencer-Milnes X, de Arroyo Garcia L, Paschou D, Lazenbatt C, Kong D, Chughtai H, Jensen K, Freemont PS, Kitney R, Reeve B, Ellis T. 2016. Engineering control of bacterial cellulose production using a genetic toolkit and a new cellulose-producing strain. Proc Natl Acad Sci U S A 113:E3431–40.

14. Geng W, Wang L, Jiang N, Cao J, Xiao Y-X, Wei H, Yetisen AK, Yang X-Y, Su B-L. 2018. Single cells in nanoshells for the functionalization of living cells. Nanoscale 10:3112–3129.

15. Gasperini L, Mano JF, Reis RL. 2014. Natural polymers for the microencapsulation of cells. J R Soc Interface 11:20140817.

16. Martín MJ, Lara-Villoslada F, Ruiz MA, Morales ME. 2015. Microencapsulation of bacteria: A review of different technologies and their impact on the probiotic effects. Innov Food Sci Emerg Technol 27:15–25.

17. Dube DH, Bertozzi CR. 2003. Metabolic oligosaccharide engineering as a tool for glycobiology. Curr Opin Chem Biol 7:616–625.

18. Ming Miao, Eva Sitarz, Catherine M. Bellingham, Emily Won, Lisa D. Muiznieks, Fred W. Keeley. 2013. Sequence and domain arrangements influence mechanical properties of elastin-like polymeric elastomers. Biopolymers 99:392–407.

19. Manoj B. Charati, Jamie L. Ifkovits, Jason A. Burdick, Jeffery G. Linhardt, Kristi L. Kiick. 2009. Hydrophilic elastomeric biomaterials based on resilin-like polypeptides. Soft Matter 5:3412–3416.

20. Linqing Li, Sean Teller, Rodney J. Clifton, Xinqiao Jia, Kristi L. Kiick. 2011. Tunable Mechanical Stability and Deformation Response of a Resilin-Based Elastomer. Biomacromolecules 12:2302–2310.

21. Nettles DL, Chilkoti A, Setton LA. 2010. Applications of elastin-like polypeptides in tissue engineering. Adv Drug Deliv Rev 62:1479–1485.

22. Muiznieks LD, Reichheld SE, Sitarz EE, Miao M, Keeley FW. 2015. Proline-poor hydrophobic domains modulate the assembly and material properties of polymeric elastin. Biopolymers 103:563–573.

23. Zakeri B, Fierer JO, Celik E, Chittock EC, Schwarz-Linek U, Moy VT, Howarth M. 2012. Peptide tag forming a rapid covalent bond to a protein, through engineering a bacterial adhesin. Proc Natl Acad Sci U S A 109:E690–7.

24. Fei Sun, Wen-Bin Zhang, Alborz Mahdavi, Frances H. Arnold, David A. Tirrell. 2014. Synthesis of bioactive protein hydrogels by genetically encoded SpyTag-SpyCatcher chemistry. Proceedings of the National Academy of Sciences 111:11269–11274.

25. Charrier M, Li D, Mann VR, Yun L, Jani S, Rad B, Cohen BE, Ashby PD, Ryan KR, Ajo-Franklin CM. 2019. Engineering the S-Layer of Caulobacter crescentus as a Foundation for Stable, High-Density, 2D Living Materials. ACS Synth Biol 8:181–190.

26. Thanbichler M, Iniesta AA, Shapiro L. 2007. A comprehensive set of plasmids for vanillate- and xylose-inducible gene expression in Caulobacter crescentus. Nucleic Acids Res 35:e137.

27. Farr C, Nomellini JF, Ailon E, Shanina I, Sangsari S, Cavacini LA, Smit J, Horwitz MS. 2013. Development of an HIV-1 microbicide based on Caulobacter crescentus: blocking infection by high-density display of virus entry inhibitors. PloS one. Public Library of Science.

28. Hillson NJ, Hu P, Andersen GL, Shapiro L. 2007. Caulobacter crescentus as a whole-cell uranium biosensor. Appl Environ Microbiol 73:7615–7621.

29. Park DM, Reed DW, Yung MC, Eslamimanesh A, Lencka MM, Anderko A, Fujita Y, Riman RE, Navrotsky A, Jiao Y. 2016. Bioadsorption of Rare Earth Elements through Cell Surface Display of Lanthanide Binding Tags. Environ Sci Technol 50:2735–2742.

30. Lasker K, Mann TH, Shapiro L. 2016. An intracellular compass spatially coordinates cell cycle modules in Caulobacter crescentus. Curr Opin Microbiol 33:131–139.

31. Tsang PH, Li G, Brun YV, Freund LB, Tang JX. 2006. Adhesion of single bacterial cells in the micronewton range. Proc Natl Acad Sci U S A 103:5764–5768.

32. Bingle WH, Nomellini JF, Smit J. 1997. Cell-surface display of a Pseudomonas aeruginosa strain K pilin peptide within the paracrystalline S-layer of Caulobacter crescentus. Mol Microbiol.

33. Hentchel KL, Reyes Ruiz LM, Curtis PD, Fiebig A, Coleman ML, Crosson S. 2019. Genome-scale fitness profile of Caulobacter crescentus grown in natural freshwater. ISME J 13:523–536.

34. Smit J, Engelhardt H, Volker S, Smith SH, Baumeister W. 1992. The S-layer of Caulobacter crescentus: three-dimensional image reconstruction and structure analysis by electron microscopy. J Bacteriol 174:6527–6538.

35. Nomellini JF, Duncan G, Dorocicz IR, Smit J. 2007. S-layer-mediated display of the immunoglobulin G-binding domain of streptococcal protein G on the surface of Caulobacter crescentus: development of an immunoactive reagent. Appl Environ Microbiol 73:3245–3253.

36. Reddington SC, Howarth M. 2015. Secrets of a covalent interaction for biomaterials and biotechnology: SpyTag and SpyCatcher. Curr Opin Chem Biol 29:94–99.

37. Azam A, Li C, Metcalf KJ, Tullman-Ercek D. 2016. Type III secretion as a generalizable strategy for the production of full-length biopolymer-forming proteins. Biotechnol Bioeng 113:2313–2320.

38. Burdette LA, Leach SA, Wong HT, Tullman-Ercek D. 2018. Developing Gram-negative bacteria for the secretion of heterologous proteins. Microb Cell Fact 17:196.

39. Awram P, Smit J. 1998. The Caulobacter crescentus paracrystalline S-layer protein is secreted by an ABC transporter (type I) secretion apparatus. J Bacteriol 180:3062–3069.

40. Bingle WH, Nomellini JF, Smit J. 2000. Secretion of the Caulobacter crescentus S-layer protein: further localization of the C-terminal secretion signal and its use for secretion of recombinant proteins. J Bacteriol 182:3298–3301.

41. Spitz O, Erenburg IN, Beer T, Kanonenberg K, Holland IB, Schmitt L. 2019. Type I Secretion Systems-One Mechanism for All? Microbiol Spectr 7.

42. Toporowski MC, Nomellini JF, Awram P, Smit J. 2004. Two outer membrane proteins are required for maximal type I secretion of the Caulobacter crescentus S-layer protein. Journal of bacteriology. Am Soc Microbiol.

43. Wade H. Bingle, John F. Nomellini, John Smit. 2000. Secretion of the Caulobacter crescentusS-Layer Protein: Further Localization of the C-Terminal Secretion Signal and Its Use for Secretion of Recombinant Proteins. J Bacteriol 182:3298–3301.

44. Cao Y, Liu D, Zhang W-B. 2017. Supercharging SpyCatcher toward an intrinsically disordered protein with stimuli-responsive chemical reactivity. Chem Commun 53:8830–8833.

45. Roberts S, Dzuricky M, Chilkoti A. 2015. Elastin-like polypeptides as models of intrinsically disordered proteins. FEBS Lett 589:2477–2486.

46. Chung C, Lampe KJ, Heilshorn SC. 2012. Tetrakis(hydroxymethyl) phosphonium chloride as a covalent cross-linking agent for cell encapsulation within protein-based hydrogels. Biomacromolecules 13:3912–3916.

47. Madl CM, Katz LM, Heilshorn SC. 2018. Tuning Bulk Hydrogel Degradation by Simultaneous Control of Proteolytic Cleavage Kinetics and Hydrogel Network Architecture. ACS Macro Lett 7:1302–1307.

48. Marks ME, Castro-Rojas CM, Teiling C, Du L, Kapatral V, Walunas TL, Crosson S. 2010. The genetic basis of laboratory adaptation in Caulobacter crescentus. J Bacteriol 192:3678–3688.

49. Shcherbo D, Murphy CS, Ermakova GV, Solovieva EA, Chepurnykh TV, Shcheglov AS, Verkhusha VV, Pletnev VZ, Hazelwood KL, Roche PM, Lukyanov S, Zaraisky AG, Davidson MW, Chudakov DM. 2009. Far-red fluorescent tags for protein imaging in living tissues. Biochem J 418:567–574.

50. Gandham L, Nomellini JF, Smit J. 2012. Evaluating secretion and surface attachment of SapA, an S-layer-associated metalloprotease of Caulobacter crescentus. Archives of microbiology. Springer.

51. Delepelaire P. 2004. Type I secretion in gram-negative bacteria. Biochim Biophys Acta 1694:149–161.

52. Debarbieux L, Wandersman C. 2001. Folded HasA inhibits its own secretion through its ABC exporter. EMBO J 20:4657–4663.

53. Nicolynn E. Davis, Sheng Ding, Ryan E. Forster, Daniel M. Pinkas, Annelise E. Barron. 2010. Modular enzymatically crosslinked protein polymer hydrogels for in situ gelation. Biomaterials 31:7288–7297.

54. Paul A Guerette, Shawn Hoon, Dawei Ding, Shahrouz Amini, Admir Masic, Vydianathan Ravi, Byrappa Venkatesh, James C Weaver, Ali Miserez. 2014. Nanoconfined β-Sheets Mechanically Reinforce the Supra-Biomolecular Network of Robust Squid Sucker Ring Teeth. ACS Nano 8:7170–7179.

55. Cormack BP, Valdivia RH, Falkow S. 1996. FACS-optimized mutants of the green fluorescent protein (GFP). Gene 173:33–38.

56. Laber JR, Dear BJ, Martins ML, Jackson DE, DiVenere A, Gollihar JD, Ellington AD, Truskett TM, Johnston KP, Maynard JA. 2017. Charge Shielding Prevents Aggregation of Supercharged GFP Variants at High Protein Concentration. Mol Pharm 14:3269–3280.

57. Valiaev A, Abu-Lail NI, Lim DW, Chilkoti A, Zauscher S. 2007.Microcantilever sensing and actuation with end-grafted stimulus-responsive elastin-like polypeptides. Langmuir 23:339–344.

58. Delepelaire P. 1998. The SecB chaperone is involved in the secretion of the Serratia marcescens HasA protein through an ABC transporter. The EMBO Journal.

59. Byun H, Park J, Kim SC, Ahn JH. 2017. A lower isoelectric point increases signal sequence–mediated secretion of recombinant proteins through a bacterial ABC transporter. J Biol Chem.

60. Schipperus R, Eggink G, de Wolf FA. 2012. Secretion of elastin-like polypeptides with different transition temperatures by Pichia pastoris. Biotechnol Prog.

61. Zajdel TJ, Baruch M, Méhes G, Stavrinidou E, Berggren M, Maharbiz MM, Simon DT, Ajo-Franklin CM. 2018. PEDOT:PSS-based Multilayer Bacterial-Composite Films for Bioelectronics. Sci Rep 8:15293.

62. Yang SH, Kang SM, Lee K-B, Chung TD, Lee H, Choi IS. 2011. Mussel-inspired encapsulation and functionalization of individual yeast cells. J Am Chem Soc 133:2795–2797.

63. John RP, Tyagi RD, Brar SK, Surampalli RY, Prévost D. 2011. Bio-encapsulation of microbial cells for targeted agricultural delivery. Crit Rev Biotechnol 31:211–226.

64. Ely B. 1991. [17] Genetics of Caulobacter crescentus, p. 372–384. In Methods in Enzymology. Academic Press.

65. Saveleva MS, Eftekhari K, Abalymov A, Douglas TEL, Volodkin D, Parakhonskiy BV, Skirtach AG. 2019. Hierarchy of Hybrid Materials-The Place of Inorganics-in-Organics in it, Their Composition and Applications. Front Chem 7:179.

66. Gu H, Ren D. 2014. Materials and surface engineering to control bacterial adhesion and biofilm formation: A review of recent advances. Frontiers of Chemical Science and Engineering 8:20–33.

67. Schmieden DT, Basalo Vázquez SJ, Sangüesa H, van der Does M, Idema T, Meyer AS. 2018. Printing of Patterned, EngineeredE. coliBiofilms with a Low-Cost 3D Printer. ACS Synthetic Biology.

68. Schaffner M, Rühs PA, Coulter F, Kilcher S, Studart AR. 2017. 3D printing of bacteria into functional complex materials. Sci Adv 3:eaao6804.

69. Huang Y, Xia A, Yang G, Jin F. 2018. Bioprinting Living Biofilms through Optogenetic Manipulation. ACS Synth Biol 7:1195–1200.

70. Asally M, Kittisopikul M, Rué P, Du Y, Hu Z, Çağatay T, Robinson AB, Lu H, Garcia-Ojalvo J, Süel GM. 2012. Localized cell death focuses mechanical forces during 3D patterning in a biofilm. Proc Natl Acad Sci U S A 109:18891–18896.

71. Huang W, Restrepo D, Jung J-Y, Su FY, Liu Z, Ritchie RO, McKittrick J, Zavattieri P, Kisailus D. 2019. Multiscale Toughening Mechanisms in Biological Materials and Bioinspired Designs. Adv Mater 31:e1901561.

72. Spiesz EM, Schmieden DT, Grande AM, Liang K, Schwiedrzik J, Natalio F, Michler J, Garcia SJ, Aubin-Tam M-E, Meyer AS. 2019. Bacterially Produced, Nacre-Inspired Composite Materials. Small 15:e1805312.

73. Smit J, Kaltashov IA, Cotter RJ, Vinogradov E, Perry MB, Haider H, Qureshi N. 2008. Structure of a novel lipid A obtained from the lipopolysaccharide of Caulobacter crescentus. Innate Immun 14:25–36.

74. Amat F, Comolli LR, Nomellini JF, Moussavi F, Downing KH, Smit J, Horowitz M. 2010. Analysis of the intact surface layer of Caulobacter crescentus by cryo-electron tomography. Journal of bacteriology. Am Soc Microbiol.

75. Persat A, Stone HA, Gitai Z. 2014. The curved shape of Caulobacter crescentus enhances surface colonization in flow. Nature communications. Nature Publishing Group.

