## Supplemental materials for "Engineering high-yield biopolymer secretion creates an extracellular protein matrix for living materials"

**SUPPLEMENTAL INFORMATION**

**SUPPLEMENTAL FIGURES**

**Supplementary Figure 1: DNA sequences for Displayer Strain**

Genome insertion of $(GGSG)_4$-SpyTag-$(GGSG)_4$ insertion with 100 base pairs up and downstream for Disp:$RsaA_{467}$-ST:
caacgccgtgaacaccacgttgacgcaagccgacgtgaccgtgaccggtaactccagcaccacggccgtgacggtcacccaaac
cgccgccgccaccgccGGAGGCTCAGGGGGAGGTTCGGGTGGCGGTTCGGGAGGAGGCTCGG
GTGCGCATATCGTAATGGTCGATGCATACAAGCCCACGAAAGGAGGTTCAGGCGGCGGAA
GCGGTGGTGGAAGCGGAGGTGGGTCAGGCggcgctacggtcgccggtcgcgtcaacggcgctgtgacgatca
ccgactctgccgccgcctcggccacgaccgccggcaagatcgccacggtcaccctgg

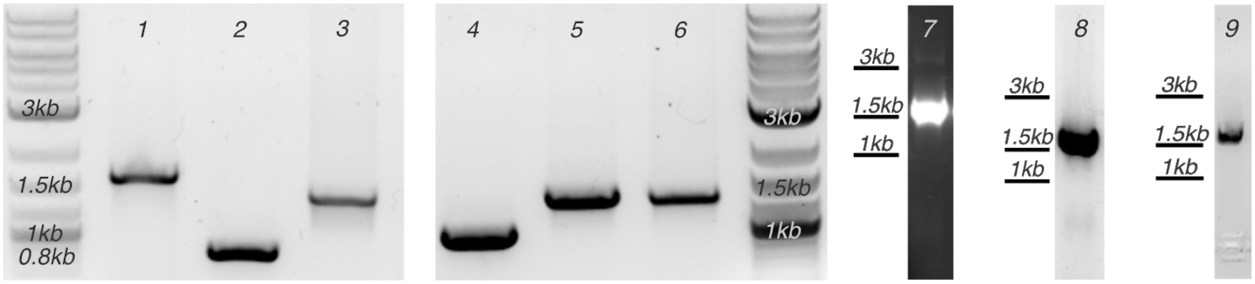


**Supplementary Figure 2: Colony PCR confirmation of genetic insertions**

Colony PCR confirm successful genomic insertion of 1) MFm127, SC-336c, 1.6kb 2) MFm142, 336c, 0.8kb 3) MFm144, SC(-)-ELP60-336c, 1.3kb 4) MFm152, ELP60-336c, 1kb 5) MFm159, SC-ELP60-336c, 1.3kb 6) MFm161, SC(-)-ELP60x-336c, 1.3kb 7) MFm149, SC(-)-RLP12-336c, 1.5kb 8) MFm151, SC(-)-Suckerin19-336c, 1.5kb 9) MFm109, RsaA467:SpyTag, 1.6kb. Molecular weight markers correspond to New England Biosciences 2-Log DNA Ladder.

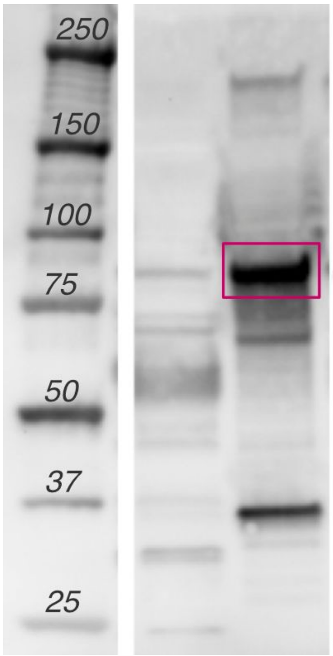


**Supplementary Figure 3: SC$^{(-)}$-ELP$_{60}$-336c is detected with anti-RsaA antibodies.**

Immunoblot with anti-RsaA polyclonal antibodies targeting purified SC$^{(-)}$-ELP$_{60}$c protein in the whole cell lysate (lane 1) and extracellular media (lane 2). Purple box indicates the full-length SC$^{(-)}$-ELP$_{60}$-336c protein. Molecular weight markers of the ladder are in kilodalton.

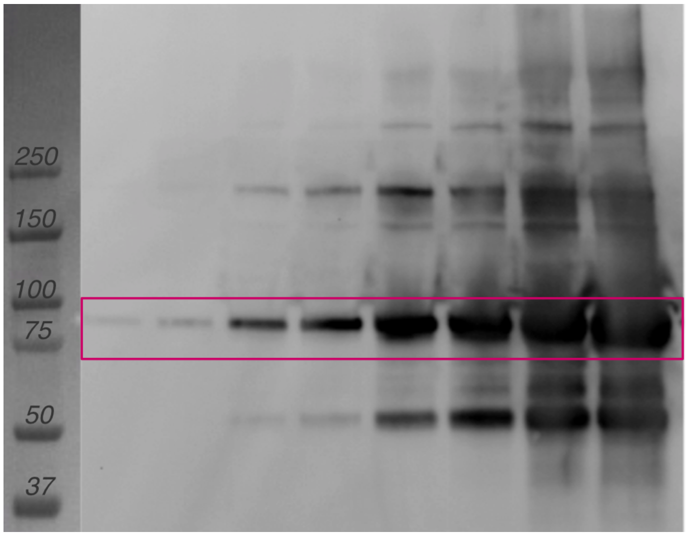


**Supplementary Figure 4: Aggregation bands increase in response to concentration of matrix protein.**

Immunoblot with anti-FLAG antibodies targeting purified SC$^{(-)}$-ELP$_{60}$-336c protein. Loading concentration is increased from left to right (0.0005, 0.001, 0.005, 0.01, 0.05, 0.1, 0.5, 1 mg/ml.)

Purple box corresponds to expected molecular weight of SC$^{(-)}$-ELP$_{60}$-336c. Molecular weight markers of the ladder are in kilodalton.

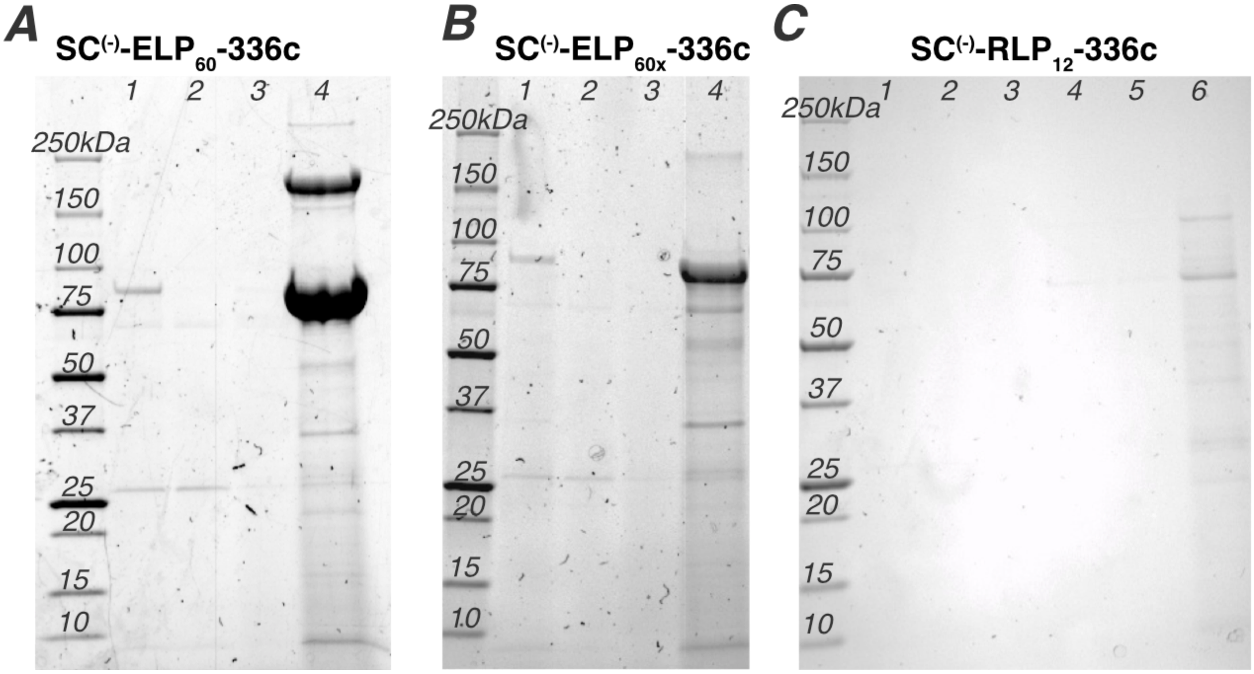


**Supplementary Figure 5: Purification of matrix proteins**

SDS-PAGE of ion exchange chromatography of extracellular media containing A) SC$^{(-)}$-ELP$_{60}$-336c B)SC$^{(-)}$-ELP$_{60x}$-336c C) SC$^{(-)}$RLP$_{12}$c. Lanes correspond to the 1) load 2) flow-through 3) wash and 4) elution. Panel C includes lanes 5) after-dialysis and 6) concentrated protein.

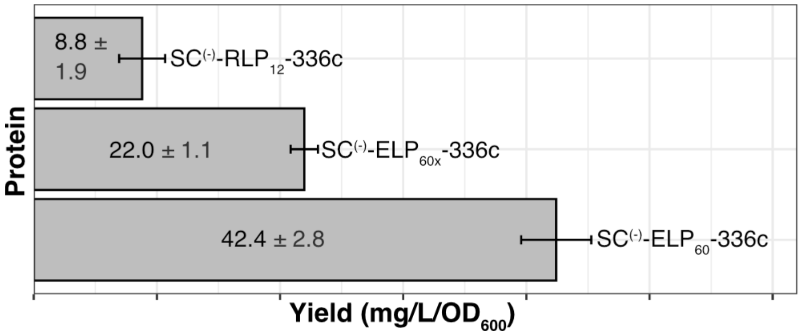


**Supplementary Figure 6: Protein concentration normalized to OD$_{600}$.**

Average yield of protein purified from the extracellular media of Sec:SC$^{(-)}$-ELP$_{60}$-336c cultures, Sec:SC$^{(-)}$-RLP$_{12}$-336c cultures, and Sec:SC$^{(-)}$-ELP$_{60x}$-336c cultures. Values in the graph are represented as average protein yield in milligrams per liter of culture per OD$_{600}$ ±SD.

### SUPPLEMENTAL METHODS

#### Plasmid construction.

For the construction of the background strains (*C. crescentus* CB15N Δ*sapA*::Pxyl-*mkate2* and *C. crescentus* CB15 Δ*sapA*::Pxyl-*GFPmut3*), a pNPTS138 integration plasmid was used to remove the S-layer protease gene (*sapA*, *CCNA_00783*) and replace it with a gene encoding a mKate2 or GFPmut2 fluorescent protein under a xylose-induction promoter. We PCR amplified 800 base pairs of the upstream and downstream regions of *sapA* using primers sapA_US and sapA_DS (Table 3) from genomic DNA purified from *C. crescentus* CB15N cells using DNEasy kit (Qiagen). The pXyl-GFPmut3 sequence (Table 4) with homology regions for assembly was synthesized by Integrated DNA Technologies. The pNPTS138 backbone was digested with HindIII-HF and SphI-HF restriction enzymes in CutSmart buffer (New England Biolabs). Subsequently, the sequence verified pNPTS138-sapA-GFP plasmid was amplified with PCR to remove *gfp* with pNPTS_sapA_F and pNPTS_sapA_R primers. The mKate2 sequence was PCR amplified with homology regions using pNPTS_mKate2-F and pNPTS_mKate2-R primers (Table 3) from the pXGFPC-2 Plac::mKate2 plasmid (75).

The p336c-SC plasmid is not used in the study except as a source plasmid for subsequent pNPTS138 construction. It was constructed from a synthetic *spycatcher* gene (Table 4, Integrated DNA Technologies) and codon-optimized for *C. crescentus.* The synthetic gene was PCR amplified with SC_336_F and SC_336_R primers and the p336c backbone plasmid (courtesy of the Smit Lab, UCB) linearized with the 336c_start_F and 336c_start_R primers.

To construct the pNPTS138-SC-336c integration plasmid, pNPTS138 was digested with NheI and HindIII-HF restriction enzymes in CutSmart buffer (New England BioLabs) for 3 hours at 37°C. The *spycatcher-336c* sequence was PCR amplified from the previously constructed p336c-SC plasmid using primers FLAG-SC-F and 336-pNPTS_R. The upstream sequence of *rsaA* (*CCNA_01059*) was amplified from the genomic DNA of *C. crescentus* CB15, and purified with a DNEasy kit (Qiagen) using US_rsaA_F and US_rsaA_R primers (Table 3). Subsequently, pNPTS138-336c was created from pNPTS138-SC-336c using the Q5 mutagenesis kit (New England BioLabs) with primers pNPTS-336_F and pNPTS_USrsaA_R. $sc^{(-)}$*-hydrogel* ($sc^{(-)}$-$elp_{60}$, $sc^{(-)}$-$rlp_{12}$, or $sc^{(-)}$-$suckerin_{19}$) fusion gene sequences were synthesized and inserted into the pUC57 plasmid (Genscript). These were digested with BamHI-HF and either ApoI-HF or EcoRI-HF restriction enzymes in CutSmart buffer (New England BioLabs) for 4-5 hours at 37°C. The digests were then assembled with a pNPTS138-SC-336c plasmid linearized to remove *sc* with the primer pairs 336-scSC-ELP60_F with pNPTS_superSC_R2, 336-scSC-RLP-F with pNPTS_superSC_R or 336-scSC-Suckerin-F with pNPTS_superSC_R. pNPTS138-SC$^{(-)}$-ELP60$_x$-336c was constructed in its entirety by Genscript. pNPTS138-ELP$_{60}$-336c was constructed from the pNPTS138-336c plasmid linearized with primers pNPTS_ELP60_F and pNPTS_ELP60_R and and $elp_{60}$ digested with Apo1-HF and BamHI-HF in CutSmart buffer for 5 hours at 37°C from the pUC57 source plasmid. pNPTS138-SC-ELP$_{60}$-336c was constructed from pNPTS138-336c linearized with primers 336c-scSC-ELP60-F and pNPTS138-336c_R, *spycatcher* amplified from p336c-SC with FLAG-SC_F and SC-STREP_R using Phusion High-Fidelity PCR Master Mix with HF Buffer, and $elp_{60}$ digested from p336c-ELP$_{60}$ with ApoI-HF and BamHI-HF at 37°C for 6 hours.

All PCR and restriction digest fragments were visualized on 1% agarose gels with 1x SYBR Safe DNA Gel stain (ThermoFisher Scientific). DNA bands were excised and purified with QX1 solubilization buffer (Qiagen) and the DNA Clean & Concentrate kit (Zymo Research) according to the manufacturer's guidelines. All PCR reactions were performed using Q5 High Fidelity 2x Master Mix (New England BioLabs) according to the manufacturer's guidelines.

Gibson Assembly was carried out using HiFi Gibson Assembly 2x Master Mix (New England BioLabs) for 1 hour at 50°C. 5 µl of each Gibson assembly reaction was transformed into NEB 5-alpha chemically competent cells (New England Biolabs) and plated on LB supplemented with kanamycin to select for the pNPTS138 plasmids. Successful assemblies were confirmed by Sanger sequencing at the UC Berkeley DNA Sequencing Facility (Berkeley, CA, USA).

**Supplemental Table 1. Strains used in this study**

| **Name** | **Characteristics** | **Plasmid** | **Strain background** | **Source** |
|---|---|---|---|---|
| MFm092 Disp:$RsaA_{wt}$ | *C. crescentus* CB15 Δ*sapA*::Pxyl-*gfpmut3* | None | CB15 | This study |
| MFm109 Disp:$RsaA_{467}$-ST | *C. crescentus* CB15 Δ*sapA*::Pxyl-*gfpmut3* *$rsaA_{467}$-spytag* | None | CB15 | This study |
| Mfm126 Sec:parent | *C. crescentus* CB15N Δ*sapA*::Pxyl-*mKate2* | None | CB15N | This study |
| Mfm 127 Sec:SC-336c | *C. crescentus* CB15N Δ*sapA*::pXyl-*mKate2* Δ*rsaA*::PrsaA-*spycatcher-336c* | None | CB15N | This study |
| Mfm 142 Sec:336c | *C. crescentus* CB15N Δ*sapA*::pXyl-*mKate2* Δ*rsaA*::PrsaA-*336c* | None | CB15N | This study |

| | | | | |
|---|---|---|---|---|
| Mfm 144 Sec: SC$^{(-)}$-ELP$_{60}$-336c | *C. crescentus* CB15N Δ*sapA*::Pxyl-*mKate2* Δ*rsaA*::PrsaA-*sc$^{(-)}$-elp$_{60}$-336c* | None | CB15N | This study |
| Mfm 145 | B5BAC p336c-*spycatcher* | p336c-SpyCatcher | B5BAC | This study |
| Mfm 149 Sec:SC$^{(-)}$-RLP$_{12}$-336c | *C. crescentus* CB15N Δ*sapA*::Pxyl-*mKate2* Δ*rsaA*::PrsaA-*sc$^{(-)}$-rlp$_{12}$-336c* | None | CB15N | This study |
| Mfm151 Sec:SC$^{(-)}$-Suckerin$_{19}$-336c | *C. crescentus* CB15N Δ*sapA::*Pxyl-*mKate2* Δ*rsaA*::PrsaA-*sc$^{(-)}$-suckerin$_{19}$-336c* | None | CB15N | This study |
| Mfm 152 Sec:ELP$_{60}$-336c | *C. crescentus* CB15N Δ*sapA*::Pxyl-*mKate2* Δ*rsaA*::PrsaA-*elp$_{60}$-336c* | None | CB15N | This study |
| Mfm 159 Sec:SC-ELP$_{60}$-336c | *C. crescentus* CB15N Δ*sapA*::Pxyl-*mKate2* Δ*rsaA*::PrsaA-*spycatcher-elp$_{60}$-336c* | None | CB15N | This study |
| Mfm 161 Sec:SC$^{(-)}$-ELP$_{60x}$-336c | *C. crescentus* CB15N Δ*sapA::*Pxyl-*mKate2* | None | CB15N | This study |

| | | | | |
|---|---|---|---|---|
| | Δ*rsaA*::PrsaA-*spycatcher*$^{(-)}$-*elp*$_{60x}$-*336c* | | | |
| Mfe 939 WM3064 | DAP auxotroph *E. coli* strain for conjugation | None | WM3064 | W. Metcalf/ UIUC |

**Supplemental Table 2. Plasmids used in this study**

| Name | Characteristic | Source |
|---|---|---|
| puc57_SuperchargedSC-ELP$_{60}$ | Storage plasmid for *spycatcher*$^{(-)}$-*elp*$_{60}$ gene. | GenScript |
| puc57_SuperchargedSC-RLP$_{12}$ | Storage plasmid for *spycatcher*$^{(-)}$-*rlp*$_{12}$ gene. | GenScript |
| puc57_SuperchargedSC-Suckerin$_{19}$ | Storage plasmid for *spycatcher*$^{(-)}$-*suckerin*$_{19}$ gene. | GenScript |
| p336c | Plasmid for expression of proteins via the *C. crescentus* B5BAC T1SS. Contains the 336c secretion signal. | John Smit, University of British Columbia |

| | | |
|---|---|---|
| p336c-SpyCatcher | Plasmid expression of original SpyCatcher with 336c signal through T1SS. | This study |
| p336c-$ELP_{60}$ | Plasmid expression of *$elp_{60}$* with *336c* signal through T1SS. | This study |
| pXGFPC-2 Plac::mKate2 | Contains *mKate2* sequence. | Persat(75) |
| pNPTS138 | Plasmid for genomic insertion using two-step recombination with SacB counterselection. | M. R. K. Alley |
| pNPTS138-ΔsapA::Pxyl-mKate2 | Genomic insertion of *mKate2* under xylose induction in place of *sapA* gene. | This study |
| pNPTS138-SC-336c | Genomic insertion of *spycatcher* in place of *$rsaA_{1-690}$*. | This study |
| pNPTS138-336c | Genomic removal of *$rsaA_{1-690}$* leaving *336c* secretion signal sequence. | This study |

| | | |
|---|---|---|
| pNPTS138-ELP$_{60}$-336c | Genomic insertion of *elp$_{60}$* in place of *rsaA$_{1-690}$*. | This study |
| pNPTS138-SC-ELP$_{60}$-336c | Genomic insertion of *spycatcher-elp$_{60}$* in place of *rsaA$_{1-690}$*. | This study |
| pNPTS138-superchargedSC-336c | Genomic insertion of *spycatcher$^{(-)}$* in place of *rsaA$_{1-690}$*. | This study |
| pNPTS138-superchargedSC-ELP$_{60}$-336c | Genomic insertion of *spycatcher$^{(-)}$-elp$_{60}$* in place of *rsaA$_{1-690}$*. | This study |
| pNPTS138-superchargedSC-RLP$_{12}$-336c | Genomic insertion of *spycatcher$^{(-)}$-rlp$_{12}$* in place of *rsaA$_{1-690}$*. | This study |
| pNPTS138-superchargedSC-Suckerin$_{19}$-336c | Genomic insertion of *spycatcher$^{(-)}$-suckerin$_{19}$* in place of *rsaA$_{1-690}$*. | This study |
| pNPTS138-superchargedSC-ELP60x-336c | Genomic insertion of *supercharged spycatcher-elp$_{60x}$* in place of *rsaA$_{1-690}$*. | This study |

**Supplemental Table 3. Primers used in this study.**

| Name | Characteristics | Sequence |
|---|---|---|
| pNPTS_sapA_R | To amplify pNPTS138-sapA-eGFP removing *eGFP* | CATGTCGTCTCCCCAAAACTCGAGCGTCTGAAGC |
| pNPTS_sapA_F | To amplify pNPTS138-sapA-eGFP, removing *eGFP* | CCGTTCGAAGGGCGCGGCGAC |
| pNPTS-mKate2_F | To amplify *mKate2* with overlaps to pNPTS138-sapA | gacgctcgagttttggggagacgacATGGTGAGCGAGCTGATTAAGGAGAACATG |
| pNPTS-mKate2_R | To amplify *mKate2* with overlaps to pNPTS138-sapA | gtcgccgcgcccttcgaacggtcatctGTGCCCCAGTTTGCTAGGGAGGTC |
| 336-pNPTS_R | To obtain *sc-336c* from p336c-SC. | cgcgtcacggccgaagctagcGTCGCAGCAGCGCCCAGGGTG |

| | | |
|---|---|---|
| FLAG-SC-F | To amplify *sc-336c* from p336c-SC. | gactacaaggacgacgatgacaagGCGATGGTGGATACGCTCTCCGGC |
| US_rsaA_F | To amplify upstream *rsaA* from CB15 genome. | caattgaagccggctggcgccaagcttCTCAGGCCGCGATCAGTGCCGAC |
| US_rsaA_R | To amplify upstream *rsaA* from CB15 genome. | catcgccttgtcatcgtcgtccttgtagtcCATGAGGATTGTCTCCCAAAAAAAATCCCACACCC |
| 336-scSC-F | To amplify pNPTS138-SC-336c with overlap to *spycatcher*$^{(-)}$. Includes *336c* sequence. Excises original *spycatcher* gene | cacatcgacgggggctgcggcaaatttGCTGACCCGGCCTTCGGCGGC |
| pNPTS_superSC_R | To amplify pNPTS138-SC-336c with overlap to *spycatcher*$^{(-)}$. | gcgtgtccaccatggccatgaattcCTTGTCATCGTCGTCCTTGTAGTCCATGAGG |
| pNPTS_superSC_R2 | To amplify pNPTS138-SC-336c with overlap to *spycatcher*$^{(-)}$. | CGGACAGCGTGTCCACCATGGCCATgaattcCTTGTCATCGTCGTCCTTGTAGTCCATGAGG |

| | | |
|---|---|---|
| 336-scSC-ELP60_F | Amplify pNPTS138-336c with overlaps to *sc-elp$_{60}$-336c*. | caggcggctcgggcggatccGCTGACCCGGCCTTCGGCGG |
| pNPTS138-336c_R | Amplify pNPTS138-336c with overlaps to *sc-elp$_{60}$-336c.* | CTTGTCATCGTCGTCCTTGTAGTCCATGAGGATTGTCTCCC |
| 336-scSC-RLP F | To amplify pNPTS138-SC-336c with overlap to *rlp$_{12}$*. Includes *336c* sequence. Excises original *spycatcher* gene. | cctggggggcggctcgggcggatccGCTGACCCGGCCTTCGGCGG |
| 336-scSC-Suckerin F | To amplify pNPTS138-SC-336c with overlap to *suckerin$_{19}$*. Includes *336c* sequence. Excises original *spycatcher* gene. | gctccacggcggctcgggcggatccGCTGACCCGGCCTTCGGCGG |
| pNPTS_USrsaA_R | To amplify pNPTS138-SC-336c with no overlap. Includes *336c* sequence. Excises original *spycatcher* gene. | CTTGTCATCGTCGTCCTTGTAGTCCATGAGGATTGTCTCCC |

| | | |
|---|---|---|
| pNPTS-336_F | To amplify pNPTS138-SC-336c with no overlap. Includes *336c* sequence. Excises original *spycatcher* gene. | GCTGACCCGGCCTTCGGCGGC |
| pNPTS-336_R | To amplify pNPTS138-SC-336c with no overlap. Includes *336c* sequence. Excises original *spycatcher* gene. | CTTTTCAAACTGCGGGTGGGACCAaaatttGCC |
| FLAG-SC_F | Amplify *spycatcher* with overlaps to backbone and *strep-elp*$_{60}$. | gactacaaggacgacgatgacaagGCGATGGTGGATACGCTCTCCGGC |
| SC-STREP_R | Amplify *spycatcher* with overlaps to backbone and *strep-elp*$_{60}$. | gcgggtgggaccaaaatttGCCGCTCCCGCCGTCGATATG |

| | | |
|---|---|---|
| pNPTS_ELP60_F | To amplify pNPTS138-336c with overlap to *elp*$_{60}$. | CCAGGCGGCTCGGGCggatccGCTGACCCGG CCTTCGGCGG |
| pNPTS_ELP60_R | To amplify pNPTS138-336c with overlap to *elp*$_{60}$. | AACTCCCTTTTCAAACTGCGGGTGGGACCAa aatttCTTGTCATCGTCGTCCTTGTAGTCCATGA GGATTG |
| SacB-F | Colony PCR primer to confirm removal of *SacB* from genome. | GGAAGCTCGGCGCAAACGTTGATTG |
| SacB-R | Colony PCR primer to confirm removal of *SacB* from genome. | CCACATCGTCTTTGCATTAGCCGGAGATCC |
| KI-336c_R | Colony PCR primer, to confirm removal of *rsaA*, leaving only *flag-336c*. | CAGCCTTGTCATCGTCGTCCTTGTAGTCC |

| | | |
|---|---|---|
| KI_RLP12_R | Colony PCR primer, to confirm knock-in of *sc*$^{(-)}$-*rlp*$_{12}$ genes after PrsaA and before *336c*. | GAGTCGGACGGACGCCCGCCATTC |
| KI_Suckerin_R | Colony PCR primer, to confirm knock-in of *sc*$^{(-)}$-*suckerin*$_{19}$ genes after PrsaA and before *336c*. | CCAGGCCGTAGCCGCCGTAGAGG |
| KI_SC_F | Colony PCR primer, knock-in confirmation of *spycatcher*. | GACGAACCAGGGTTCGTTCTCGTCGC |
| ColonyPCR_336c_R | Colony PCR primer, knock-in confirmation of 336c. | GATCGACTTGGCCGAGGTGGCTTGCA |
| ColonyPCR_ELP60_R | Colony PCR primer, knock-in confirmation of *sc*$^{(-)}$-*elp*$_{60}$ and *elp*$_{60}$. | ACACCTGCTCCGGGAACTCCCCCA |
| ColonyPCR_ELP60X_R | Colony PCR primer, knock-in confirmation of *sc*$^{(-)}$-*elp*$_{60x}$. | CCCTTGCTGGCCTGGAACTCCTA |

| | | |
|---|---|---|
| SC_336_F | To amplify the *ggsg_spycatcher_CB* gene block | caatttcacacaggaaacagctatgGCGATGGTGGATAC GCTCTCCGGCCTGTCG |
| SC_336_R | To amplify the *ggsg_spycatcher_CB* gene block | cgggaattcgtaatcatggtGCCGCTCCCGCCGTCGAT ATGGGCGTCGCCCTTGGTCGCC |
| 336C_start_F | To linearize the p336c plasmid. | ACCATGATTACGAATTCCCGGGGATCC |
| 336C_start_R | To linearize the p336c plasmid. | CATAGCTGTTTCCTGTGTGAAATTGTTATCCGC |

**Supplemental Table 4. Synthesized gene blocks used in this study.**

| **Name** | **Characteristics** | **Sequence** |
|---|---|---|
| GGSG_SpyCatcher_CB | SpyCatcher with GGSG linker sequence codon optimized for *C. crescentus*. Synthesized by Integrated DNA Technologies | GCGATGGTGGATACGCTCTC CGGCCTGTCGTCGGAGCAAG GGCAGTCGGGGGATATGACC ATCGAGGAGGATTCGGCCAC CCACATCAAGTTCTCCAAGC GTGATGAAGACGGGAAGGAA CTCGCCGGGGCCACGATGGA GCTCCGCGACTCGTCCGGGA AGACCATCTCCACCTGGATCT CGGATGGGCAAGTGAAGGAC TTTTATCTCTACCCCGGGAAG TATACGTTTGTCGAGACGGC GGCCCCCGATGGGTACGAG GTGGCGACCGCGATCACGTT TACCGTCAATGAACAGGGGC AGGTCACCGTGAACGGCAAG GCGACCAAGGGCGACGCCC ATATCGACGGCGGGAGCGGC |

| | | |
|---|---|---|
| pXyl-GFPmut3 | pXyl-GFPmut3 sequence codon optimized for *C. crescentus.* Synthesized by Integrated DNA Technologies | ATACTCCTTTCAGGTGAGTGG<br>AGCGCGTCGCTGCAGCCAGC<br>CGTGGTCGGGCAGCAGGTAG<br>AAGGCGCCCTCGTCCTGATC<br>CTCGCCCGAAACCTCCAGCC<br>CCCGGTCGATGGCTTCGACG<br>ACATAGCCGGCCGCGCGGCA<br>GGTGTCGGTGAGCGCGGCCA<br>GCAGGGCGGCTTCCTGGTCA<br>GGGGTCAGGTCGGTCATGGG<br>CAAGAGGTCCAGGTCGTGGT<br>TTGTCGGCGGCTTCTAGCAT<br>GGACCGCCCGCGCCCGTGA<br>GGCCGAGGATTTCGCGCTGG<br>TCAGACAACCTACTTGCCGTC<br>CCCACATGTTAGCGCTACCA<br>AGTGCCGACGAACGCGCGCC<br>GCCGACGGTGTCGGCGCTTC<br>AGACGCTCGAGTTTTGGGGA<br>GACGACATGCGGAAGGGCGA<br>AGAACTCTTTACCGGGGTGG<br>TCCCGATCCTCGTGGAGCTC<br>GATGGCGATGTCAATGGGCA<br>CAAGTTCTCCGTCTCGGGCG<br>AAGGGGAGGGGGATGCCAC<br>GTATGGCAAGCTCACGCTGA<br>AGTTCATCTGCACCACCGGC<br>AAGCTCCCCGTGCCCTGGCC<br>CACCCTCGTCACGACGTTCG<br>GCTATGGCGTGCAATGCTTT<br>GCCCGTTATCCGGATCATATG<br>AAGCGCCATGATTTTTTTAAG<br>AGCGCCATGCCCGAAGGGTA<br>TGTGCAGGAACGGACCATCT<br>TCTTCAAGGACGACGGGAAT<br>TACAAGACCCGGGCGGAGGT<br>GAAGTTCGAGGGGGATACGC<br>TGGTGAATCGCATCGAGCTC<br>AAGGGGATCGACTTCAAGGA<br>GGACGGCAACATCCTGGGGC<br>ACAAGCTCGAATATAACTATA<br>ACTCGCACAATGTCTATATCA<br>TGGCCGATAAGCAAAAGAAC<br>GGGATCAAGGTGAACTTCAA<br>GATCCGGCATAACATCGAGG<br>ACGGGTCGGTGCAGCTCGCC<br>GACCATTACCAGCAGAATAC<br>GCCCATCGGGGATGGGCCG<br>GTGCTGCTGCCCGACAATCA<br>CTATCTCAGCACGCAATCCG<br>CGCTGTCGAAGGACCCGAAT<br>GAAAAGCGCGACCATATGGT<br>CCTGCTGGAGTTTGTCACGG |

| | | |
|---|---|---|
| | | CCGCCGGCATCACGCATGGC ATGGATGAACTGTATAAGTAA CCGTTCGAAGGGCGCGGCG ACAAAGGTCCA |

**Supplementary Table 5: Isoelectric points of extracellular matrix proteins and individual protein domains.**

| Protein/Domain | Isoelectric point (pI) |
|---|---|
| 336c | 3.83 |
| SC | 4.46 |
| SC-336c | 4.14 |
| SC-$ELP_{60}$-336C | 4.16 |
| $ELP_{60}$-336c | 3.99 |
| $SC^{(-)}$ | 3.98 |
| $SC^{(-)}$-$ELP_{60}$-336c | 3.97 |
| $SC^{(-)}$-$RLP_{12}$-336c | 4.33 |
| $SC^{(-)}$-$ELP_{60x}$-336c | 5.09 |
| $ELP_{60}$ | 5.52 |
| $RLP_{12}$ | 9.91 |
| $ELP_{60x}$ | 10.7 |
| $Suckerin_{19}$ | 8.33 |
